# Dual-targeting CRISPR-CasRx reduces *C9orf72* ALS/FTD sense and antisense repeat RNAs in vitro and in vivo

**DOI:** 10.1101/2024.01.26.577366

**Authors:** Liam Kempthorne, Deniz Vaizoglu, Alexander J. Cammack, Mireia Carcolé, Pacharaporn Suklai, Bhavana Muralidharan, François Kroll, Thomas G. Moens, Lidia Yshii, Stijn Verschoren, Benedikt V. Hölbling, Eszter Katona, Alla Mikheenko, Rachel Coneys, Paula de Oliveira, Yong-Jie Zhang, Karen Jansen, Lillian M Daughrity, Alexander McGown, Tennore M. Ramesh, Ludo Van Den Bosch, Ahad A. Rahim, Leonard Petrucelli, Jason Rihel, Adrian M. Isaacs

## Abstract

The most common genetic cause of both frontotemporal dementia (FTD) and amyotrophic lateral sclerosis (ALS) is a G_4_C_2_ repeat expansion in intron 1 of the *C9orf72* gene. This repeat expansion undergoes bidirectional transcription to produce sense and antisense repeat RNA species. Both sense and antisense-derived repeat RNAs undergo repeat-associated non-AUG translation in all reading frames to generate five distinct dipeptide repeat proteins (DPRs). Importantly, toxicity has been associated with both sense and antisense repeat-derived RNA and DPRs. This suggests targeting both sense and antisense repeat RNA may provide the most effective therapeutic strategy. The RNA-targeting CRISPR-Cas13 systems offer a promising avenue for simultaneous targeting of multiple RNA transcripts, as they mature their own guide arrays, thus allowing targeting of more than one RNA species from a single construct. We show that CRISPR-Cas13d originating from *Ruminococcus flavefaciens* (CasRx) can successfully reduce *C9orf72* sense and antisense repeat transcripts and DPRs to background levels in HEK cells overexpressing *C9orf72* repeats. CRISPR-CasRx also markedly reduced the endogenous sense and antisense repeat RNAs and DPRs in three independent *C9orf72* patient-derived iPSC-neuron lines, without detectable off-target effects. To determine whether CRISPR-CasRx is effective *in vivo*, we treated two distinct *C9orf72* repeat mouse models using AAV delivery and observed a significant reduction in both sense and antisense repeat-containing transcripts. Taken together this work highlights the potential for RNA-targeting CRISPR systems as therapeutics for *C9orf72* ALS/FTD.

## Introduction

Amyotrophic lateral sclerosis (ALS) and frontotemporal dementia (FTD) are two inexorable neurodegenerative disorders that exist on a common disease spectrum, characterised by motor impairment and behavioral and language deficits, respectively ^1^. The most common genetic cause of both ALS and FTD is a G_4_C_2_ hexanucleotide repeat expansion in intron 1 of chromosome 9 open reading frame 72 (*C9orf72*) ^2,3^. Healthy individuals typically harbour up to 30 G_4_C_2_ repeats, whereas those with *C9orf72*-related ALS and FTD (C9 ALS/FTD) may have hundreds to thousands of G_4_C_2_ repeats ^4^. These repeat-containing regions are bidirectionally transcribed into both sense (G_4_C_2_) and antisense (C_4_G_2_) repeat-containing transcripts ^5–8^. Both transcripts undergo repeat-associated non-AUG (RAN) translation, leading to the production of five distinct species of dipeptide repeat proteins (DPRs): polyGA, polyGP and polyGR from the sense strand and polyGP, polyPA and polyPR from the antisense strand ^6,7,9–11^.

To date, therapeutic strategies to target C9 ALS/FTD repeat RNAs have focused on targeting the sense G_4_C_2_ repeat transcripts ^12–16^, however, there is a considerable and growing body of evidence that supports the contribution of the antisense strand to C9 ALS/FTD disease pathogenesis ^17–21^. Additionally, recent failures in clinical trial of two antisense oligonucleotides (ASOs) targeting *C9orf72* sense repeat-containing transcripts (BIIB078; clinicaltrials.gov: NCT03626012 and WVE-004; clinicaltrials.gov: NCT04931862), despite showing target engagement and lowering of the sense transcript, highlight the importance of developing therapeutic approaches that can also target *C9orf72* antisense repeat RNA transcripts.

Clustered regularly interspaced short palindromic repeat (CRISPR) RNAs and CRISPR-associated (Cas) proteins are part of the adaptive immunity of bacteria ^22^. The recent discovery of Type VI Cas13 effectors with ribonuclease activity represent a breakthrough in RNA interference systems ^23–26^. Cas13 effectors have been shown to exhibit dual ribonuclease activity in mammalian systems, with the ability to mature their own guide RNA (gRNA) arrays from a CRISPR locus as well as degrade targeted RNA strands via higher eukaryotes and prokaryotes nucleotide-binding (HEPN) domains ^27^. This dual ribonuclease activity, combined with the smaller size of some Cas13 effectors (such as *Rfx*Cas13d at 967 amino acids) ^26^, suggest CRISPR-Cas13 systems could be utilized as therapeutics that can be packaged into an adeno-associated virus (AAV) vector and targeted to multiple human RNA sequences ^28–30^. We therefore aimed to exploit CRISPR-Cas13 systems to target both the sense and antisense repeat RNA species in *C9orf72* ALS/FTD in a single therapeutic design.

## Results

### CRISPR-Cas13d(Rx) reduces *C9orf72* sense and antisense repeat RNA and DPRs in HEK cells

To determine whether CRISPR-Cas13 systems could efficiently degrade *C9orf72* repeat RNAs, we designed 30 nucleotide (nt) gRNAs to target sense G_4_C_2_ repeat-containing transcripts. We tiled the guides every 10 nucleotides directly upstream of the G_4_C_2_ repeat (**Fig. 1A**). Of these guides we selected those with the lowest homology to other sequences in the human transcriptome and lowest RNA secondary structure scores, to provide 6 gRNAs to take forward for experimental analysis. We focused on the region upstream of the repeat, rather than the G_4_C_2_ sequence itself due to the very high GC content of the repeat sequence and the presence of G_4_C_2_ repeats elsewhere in the genome. In addition, *C9orf72* has three transcript variants, with the repeat expansion being present in the pre-mRNA of variants 1 and 3, but within the promoter of variant 2, thereby excluding the repeat from variant 2 pre-mRNA (**Fig. 1A**). Targeting gRNAs upstream of the G_4_C_2_ repeat in intron 1 allows us to target only the repeat-containing transcripts (variants 1 and 3) whilst sparing variant 2 pre-mRNA (**Fig. 1A**).

**Figure 1.**
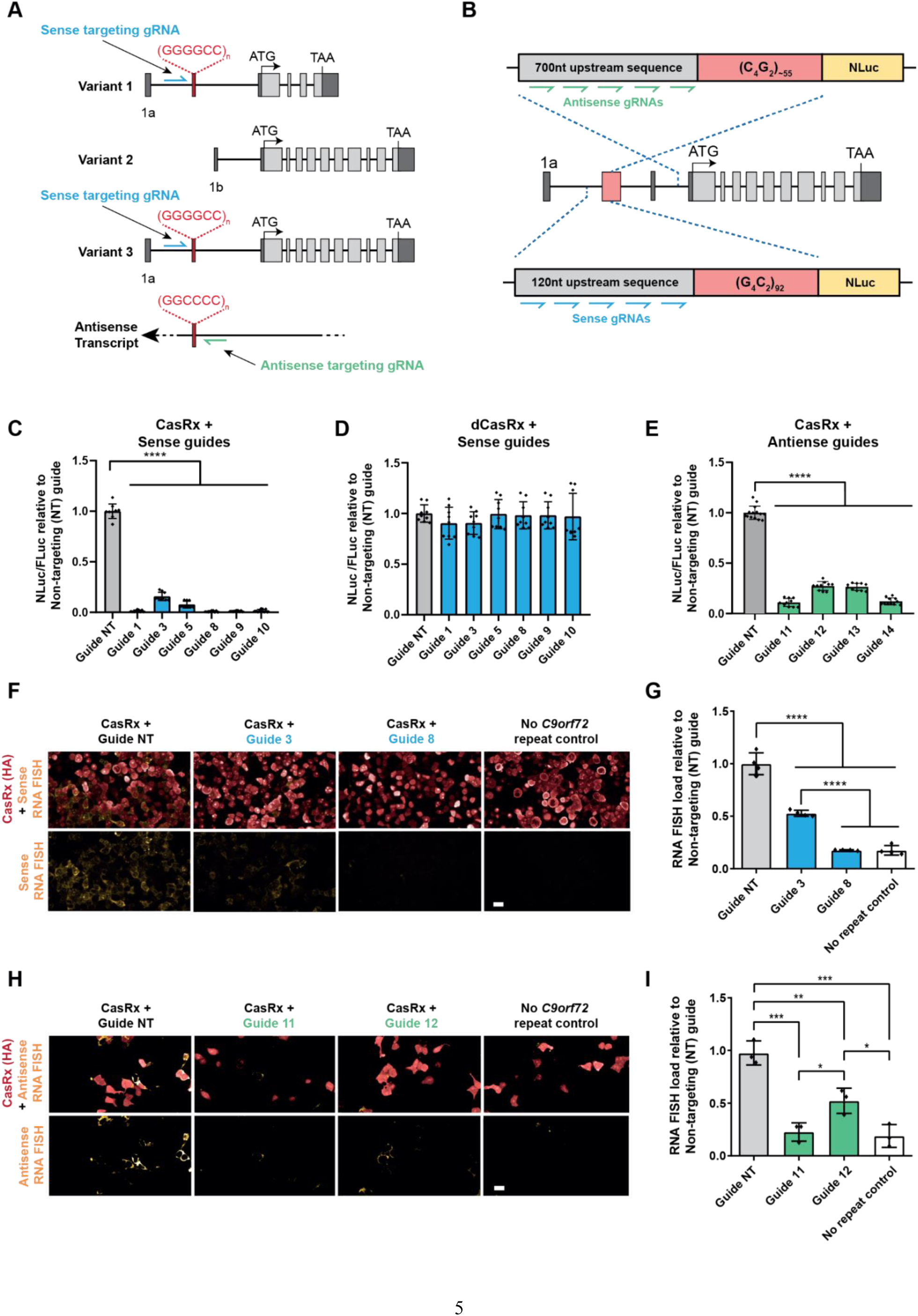
CRISPR-CasRx can effectively lower sense and antisense repeat-containing transcripts and DPRs and prevent polyGR and polyPR formation in HEK293T cells. (A) Strategy for targeting sense and antisense *C9orf72* transcripts with gRNAs. Sense gRNAs target repeat-containing variants 1 and 3 but not variant 2. (B) Sense and antisense NLuc reporter assay designs. Sense NLuc reporter assays for (C) CasRx and (D) dCasRx using sense targeting gRNAs. (E) Antisense NLuc reporter signal following CasRx and antisense targeting gRNA application. (C-E) Each NLuc reading was normalised to FLuc for each well and further normalised to the non-targeting control gRNA. Data given as mean ± S.D, n=3 biological repeats (with technical replicates shown on graph), one-way ANOVA, Holm-Sidak post-hoc analysis, ****p<0.001. (F) Representative images of RNA-FISH for the sense G_4_C_2_ transcript and ICC for the HA tag of CasRx with different CasRx guides. Scale bars = 20 µm. (G) Quantification of sense RNA FISH load calculated as integrated intensity of nuclear RNA puncta per CasRx positive cell. Data given as mean ± S.D, n=3 biological replicates, one-way ANOVA, Holm-Sidak post-hoc analysis, ****p<0.0001. (H) Representative images of RNA-FISH for the antisense C_4_G_2_ transcript and ICC for the HA tag of CasRx with different CasRx guides. Scale bars = 20 µm. (I) Quantification of antisense RNA FISH load calculated as integrated intensity of nuclear RNA puncta per CasRx positive cell. No repeat control indicates background signal of the LNA probe used for RNA FISH. Data given as mean ± S.D, n=3 biological replicates, one-way ANOVA, Holm-Sidak post-hoc analysis,*p<0.05, **p<0.01, ***p<0.001.

To test these gRNAs we utilized a NanoLuciferase (NLuc) reporter assay that we previously developed to measure sense *C9orf7*2 repeat levels (**Fig. 1B**) ^31^. Our sense NLuc reporter consists of 92 pure G_4_C_2_ repeats with 120 nucleotides of the endogenous *C9orf72* upstream sequence followed by NLuc in frame with polyGR. As the NLuc reporter has no ATG start codon, the NLuc signal is a readout of RAN-translated polyGR. We first tested two Cas13 subtypes previously shown to efficiently target mammalian transcripts, Cas13b from *Prevotella sp. P5-125* ^24^ and Cas13d from *Ruminococcus flavefaciens* strain XPD3002 (CasRx) ^25^. While Cas13b could effectively target the sense repeat-containing transcripts, with guide 8 being the most effective with a 75% (± 3% S.D.) reduction in NLuc signal (**Fig. S1A**), CasRx achieved a 99% (± 1% S.D.) reduction in NLuc signal with four of the six guides tested (guides 1, 8, 9, and 10) (**Fig. 1C)**. These data show that CasRx is more efficient than Cas13b at reducing *C9orf72* sense repeat transcripts. Importantly, in addition to the superior targeting efficiency, CasRx is 160 amino acids smaller than Cas13b, and thus is small enough to facilitate packaging, along with a gRNA array, into adeno-associated viruses (AAVs). This striking effectiveness of CasRx to target the sense repeat-containing transcripts, combined with its smaller size, led us to move forward with CasRx. We additionally tested an enzymatically dead CasRx (dCasRx) that binds to the target transcript but lacks ribonuclease activity to degrade it ^25^, as we hypothesised that binding of CasRx to repeat-containing transcripts may be sufficient to prevent translation of the repeat sequence. However, dCasRx did not reduce NLuc levels, suggesting the binding of dCasRx to the transcript is not sufficient to impair RAN translation (**Fig. 1D**). These data show that CasRx is extremely effective at reducing over-expressed *C9orf72* sense repeat RNAs, and that its ribonuclease activity is required to prevent DPR production.

As CRISPR-CasRx was very effective at reducing *C9orf72* sense repeat RNAs, we next tested whether it could also target antisense repeat-containing transcripts. We first developed an antisense NLuc reporter consisting of ∼55 pure C_4_G_2_ repeats and 700 nucleotides of the endogenous *C9orf72* sequence upstream of the repeat expansion in the antisense strand with NLuc in frame with polyPR (**Fig. 1B**). We tested 4 gRNAs targeting upstream of the antisense C_4_G_2_ in our antisense NLuc assay. These initial gRNAs were targeted as close to the C_4_G_2_ repeats as possible due to the unknown antisense transcriptional start site. All tested guides reduced the NLuc signal >70% with guide 11 achieving the greatest reduction of 89% (±4% S.D.) (**Fig. 1E**) despite the high GC-content (∼80%) of the targeted region adjacent to the antisense repeats. Finally, in a parallel approach, we used repeat-targeted RNA fluorescent *in situ* hybridization (FISH) to directly confirm *C9orf72* transcript lowering. Indeed, both sense and antisense targeted gRNAs were able to reduce *C9orf72* repeat RNA levels to near-background levels (**Fig. 1F-I**). Thus, CasRx is able to robustly degrade sense and antisense *C9orf72* repeat RNAs and prevent DPR accumulation *in vitro*.

### Dual targeting CRISPR-CasRx effectively reduces *C9orf72* sense and antisense repeat RNAs simultaneously in HEK cells

In these preceding experiments our gRNAs were expressed without the need for CasRx to mature a pre-gRNA prior to target recognition and binding. As the gRNA in a therapeutic vector would be in a pre-gRNA configuration, we next investigated whether CasRx could mature a pre-gRNA and maintain targeting efficiency. We cloned our gRNAs into a pre-gRNA expression vector where the gRNA spacer sequence is flanked by two direct repeats (DRs) and therefore must be matured prior to target engagement. Testing these pre-gRNAs in our sense NLuc assay confirmed that CasRx can mature a pre-gRNA and still very effectively target the sense repeat-containing transcripts (**Fig. S1B**). In addition, when CasRx matures a pre-gRNA array, approximately 8 nt of the guide sequence can be removed along with 6 nt of the DR to form a mature gRNA. Therefore, we also tested 22 nt gRNA versions of our most efficacious 30 nt gRNAs, with 8 nt removed from the 3’ end of the guide sequence. In our NLuc reporter assays these 22 nt gRNAs successfully targeted the sense and antisense repeat-containing transcripts (**Fig. S1C-D**).

We then cloned our most efficacious sense and antisense-targeting gRNAs into a single lentiviral vector also expressing CasRx. In our initial NLuc reporter assays (**Fig. 1C-E**) the gRNAs were supplied in 5:1 molar ratio to CasRx. These single gRNA-CasRx expressing plasmids show that a 1:1 ratio of gRNA to CasRx is still sufficient to effectively target the *C9orf72* sense and antisense repeat-containing transcripts (**Fig. S1E-F**). Here we also tested a new antisense-targeting gRNA (guide 17) that targets further from the repeat sequence (<200 bp upstream of the antisense repeats, where the sequence is less GC-rich) than the other antisense guides previously tested. We found guide 17 to be the most efficacious gRNA in our antisense NLuc assay, reducing polyPR-NLuc to background levels (**Fig. S1F**). We therefore used this antisense-targeting gRNA moving forward.

Having established our best performing sense and antisense targeting guides, we next produced a dual targeting lentiviral construct containing CasRx and sense targeting gRNA 10 and antisense targeting gRNA 17 in a pre-gRNA array. We then compared this dual-targeting construct to our single guide constructs in our sense and antisense NLuc reporter assays. The dual gRNA expressing plasmid effectively reduced polyGR-NLuc and polyPR-NLuc levels to a comparable degree to plasmids expressing individual mature gRNAs (**Fig. S1F-G**). These data show effective gRNA array maturation and multi-target engagement of CasRx with sense and antisense *C9orf72* repeat RNAs in HEK cells.

### CRISPR-CasRx effectively targets sense and antisense *C9orf72* repeat RNAs in patient-derived iPSC-neurons

Following the robust targeting efficiency of CasRx in our HEK293T transient overexpression model, we wanted to determine whether these gRNAs and CasRx can successfully target endogenous *C9orf72* sense and antisense repeat transcripts in *C9orf72* ALS/FTD patient-derived iPSC-neurons. We used piggyBac-mediated integration to insert a doxycycline (dox)-inducible NGN2 cassette into iPSCs for three independent *C9orf72* patient lines to allow rapid differentiation into cortical-like neurons ^32,33^. We confirmed that after 5 days of dox treatment these cells expressed neuronal markers using immunocytochemistry and RT-qPCR (**Fig. S2).** We transduced these dox-inducible *C9orf72* patient NGN2 iPSC lines with our CRISPR-CasRx and gRNA expressing lentiviruses. Transduced neuronal cells were isolated with FACS after 5 days of differentiation for target engagement assessment (**Fig. 2A**). Our sense targeting gRNAs 8 and 10 significantly reduced sense repeat transcripts by ∼30-80%, dependent on the cell line used, relative to the non-targeting gRNA control (**Fig. 2B** and **Fig. S3A**). As predicted, there was no reduction in *C9orf72* variant 2, which does not contain the repeats within its pre-mRNA (**Fig. 2C**). We then tested our most efficacious antisense targeting gRNA, guide 17. CasRx and guide 17 successfully reduced the antisense repeat-containing transcripts in all three *C9orf72* ALS/FTD iPSC-neuron lines by ∼50-95%, dependent on the cell line used (**Fig. 2D** and **Fig. S3A**). We also performed RT-qPCR analysis for CasRx expression in these samples (**Fig. S3B**) and found that CasRx expression correlated with antisense transcript knockdown efficiency: CasRx levels were highest in C9 line 2, the line with the greatest reduction of antisense repeat transcripts. Finally, we measured the levels of the DPRs polyGP and polyGA using quantitative MSD immunoassays and observed a ∼60-70% reduction in both polyGA (**Fig. 2E**) and polyGP (**Fig. 2F**) in all 3 lines, with both guides achieving similar efficacy. We are unable to detect DPRs specific to the antisense strand in these cells using our current immunoassays so it was not possible to elucidate the level of reduction in antisense DPRs. Taken together, these results show CasRx can achieve robust reduction in endogenous sense and antisense repeat RNAs across multiple patient iPSC-neuron lines.

**Figure 2.**
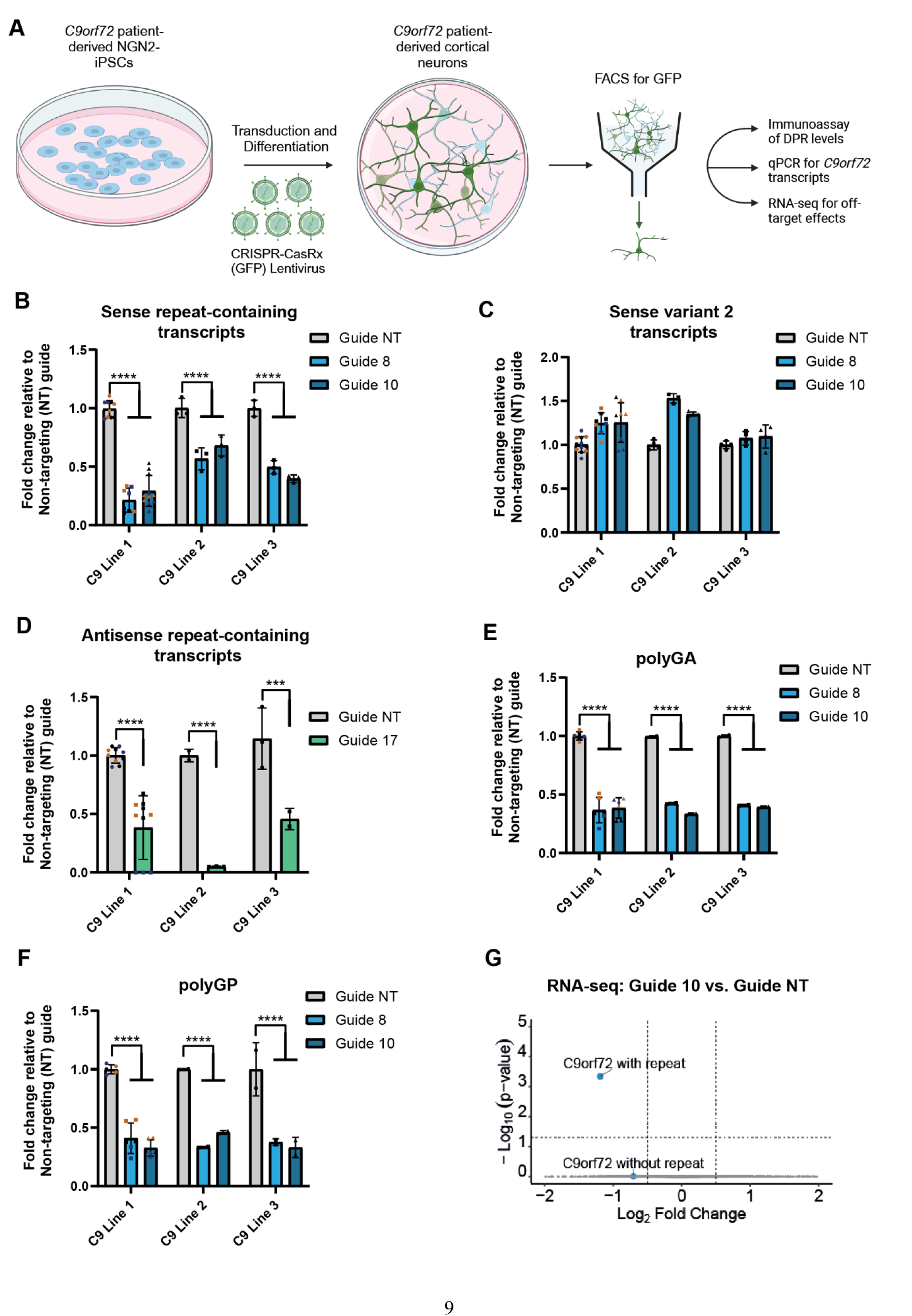
CRISPR-CasRx reduces endogenous sense and antisense repeat-containing *C9orf72* RNA and DPRs in patient-derived iPSC-neurons without lowering variant 2 or causing detectable off-target effects. **(A)** Experimental setup to determine target engagement in patient-derived iPSC-neurons. **(B-D)** RT-qPCRs for *C9orf72* transcripts in neurons treated with CRISPR-CasRx + gRNA lentiviruses. Experiments were performed on three separate inductions of C9 line 1 and in one induction each of C9 lines 2 and 3. **(B)** RT-qPCR for sense repeat-containing *C9orf72* transcripts (variants 1 and 3). **(C)** RT-qPCR for sense variant 2 *C9orf72* transcripts. **(D)** RT-qPCR for antisense repeat-containing *C9orf72* transcripts. RT-qPCR data presented as fold change compared to non-targeting (NT) gRNA. Levels of **(E)** polyGA and **(F)** polyGP DPRs in iPSC-neurons treated with sense targeting gRNAs 8 and 10, shown as fold-change compared to non-targeting (NT) gRNA. **(B-F)** Data given as mean ± S.D, n=3 biological replicates of C9 line 1 (with technical replicates) and n=3 technical replicates of C9 lines 2 and 3, each biological replicate shown as different colour data points (black, blue, and orange represent the 3 separate inductions), one-way ANOVA, Holm-Sidak post-hoc analysis, ****p<0.0001. **(G)** Volcano plot of DESeq2 analysis showing DEGs between neurons treated with sense-targeting guide 10 compared to non-targeting (NT) gRNA with *C9orf72* transcripts grouped by those that contain intron1 and the repeats, and those that do not. Dotted lines indicate thresholds for fold change on x axis (|log_2_FoldChange|>0.5) and *p* value on y-axis (adjusted p<0.05). n=3 independent inductions of C9 line 1.

### *C9orf72* repeat targeting CRISPR-CasRx does not cause toxicity or measurable off-targets in patient iPSC-neurons

There is clear evidence that activated CRISPR-Cas13 enzymes can cause toxicity due to bystander RNA cleavage when targeting highly expressed transcripts but does not have detectable off-target transcriptional changes or consequent toxicity when targeting low expression transcripts ^34^. To determine if there was overt toxicity or off-target transcriptional changes in patient iPSC-neurons when targeting *C9orf72* with CRISPR-CasRx, we performed cell viability and bulk RNA-seq analyses on one C9 patient line (C9 line 1) 5 days post-transduction. We observed no reduction in viable cells after a 5-day treatment with CasRx plus gRNA expressing lentiviruses (**Fig. S3C-D**), while a control GFP expressing lentivirus was sufficient to cause detectable levels of toxicity after 5 days (**Fig. S3D**).

Using the same experimental paradigm, we performed RNA sequencing of three independent inductions of C9 line 1 iPSC-neurons 5 days post-transduction with lentiviruses expressing CasRx and either non-targeting guide, sense transcript targeting guides 8 or 10 (**Fig. 2F** and **Fig. S3E**), or antisense transcript targeting guides 11 or 17 (**Fig. S3F-G**). For analysis of target engagement of the sense *C9orf72* transcripts we grouped the *C9orf72* transcripts into two groups as with our RT-qPCR dataset: those transcripts that contain intron 1 and the G_4_C_2_ repeat sequence (variants 1 and 3), and those that do not contain the repeat sequence (variant 2).

Guides 8 and 10 significantly reduced repeat-containing *C9orf72* transcripts but not *C9orf72* transcripts that do not contain the repeat expansion (**Fig. 2F** and **Fig. S3E**). No other differentially expressed genes (DEGs) were observed, indicating no significant off-target transcriptional changes. We were not able to detect the antisense repeat-containing transcripts in our RNA-sequencing dataset, likely due to the relatively low number of transcripts and so were not able to confirm target engagement using RNA-seq (**Fig. S3F-G**). However, our more sensitive RT-qPCR for the repeat-containing antisense transcripts did show significant reduction when targeted with CRISPR-CasRx and guide 17 (**Fig. 2D**). We did not observe any DEGs with guide 11 or guide 17 (**Fig. S3F-G**), indicating that they did not lead to off-target transcriptional changes. These data show that after 5 days of treatment in patient iPSC-neurons, *C9orf72* repeat-targeting CRISPR-CasRx does not cause overt toxicity or off-target effects.

### CRISPR-CasRx rescues larval hyperactivity in a *C9orf72* repeat zebrafish model

Having established efficacy *in vitro*, we tested CRISPR-CasRx *in vivo* using *Tg(ubi:G_4_C_2_×45)* zebrafish larvae (**Fig S4A**), which express 45 pure G_4_C_2_ repeats and generate DPRs ^35^. Using an MSD immunoassay, we confirmed that G_4_C_2_ larvae produce polyGP (**Fig. S4B**). To determine whether the model has a larval phenotype, we video-tracked two clutches of G_4_C_2_ larvae over multiple day-night cycles and assessed activity with our previously established automated pipeline ^36,37^. Compared to wild-type siblings, G_4_C_2_ larvae were markedly hyperactive (**Fig. S4C**), spending on average 1.7× more time performing swimming bouts each day (**Fig. S4D**). As this model does not contain any of the endogenous *C9orf72* human sequence upstream or downstream of the repeats, we designed new gRNAs specifically to target the zebrafish G_4_C_2_ transcripts (**Fig. S4E**). We injected single-cell embryos with plasmids encoding CasRx and G_4_C_2_-targeting or control non-targeting gRNAs, together with Tol2 recombinase mRNA to integrate the construct in the genome. We screened larvae for high expression of the construct (**Fig. S4F**) and video-tracked them (**Fig. S4H**). As expected, control-injected G_4_C_2_ larvae were hyperactive, spending 1.3× more time active each day compared to wild-type larvae expressing CasRx and non-targeting gRNAs (**Fig. S4I**). In contrast, expression of CasRx and targeting gRNAs reduced the amount of polyGP DPRs (**Fig. S4G**) and hyperactivity (**Fig. S4I**). CRISPR-CasRx can therefore rescue a behavioural phenotype of an animal model of pathogenic *C9orf72* repeats.

### CRISPR-CasRx effectively reduces sense repeat RNA in (G_4_C_2_)_149_ repeat mice

We next wanted to determine whether we could use CasRx to target sense repeats in a mouse model. We produced two PHP.eB serotype AAVs expressing CasRx and either our targeting gRNAs (guides 10 and 17), or non-targeting gRNAs as a control. We first used a mouse model that utilises neonatal intracerebroventricular (ICV) injection of an AAV expressing 149 pure G_4_C_2_ repeats (149R) ^38^. These mice have been previously shown to produce pathological hallmarks of C9 ALS/FTD including repeat-containing transcripts and DPRs ^38^. Additionally, this model contains 119 nucleotides of the endogenous *C9orf72* sequence upstream and 100 nucleotides downstream of the repeats, which means our sense-targeting gRNA (guide 10) will target the sense repeat-containing transcript in this model. However, as our antisense-targeting gRNA (guide 17) targets further from the repeats than the sequence included in this model, it cannot be used to assess our antisense guide. Postnatal day 0 (P0) mice were co-injected via ICV administration with the (G_4_C_2_)_149_ AAV and either our targeting or non-targeting CasRx PHP.eB AAVs (**Fig. 3A**). After 3 weeks we assessed levels of the sense repeat-containing transcripts in the hippocampus via RT-qPCR. This revealed a >50% reduction in sense repeat-containing transcripts, confirming successful target engagement *in vivo* (**Fig. 3B**). PolyGP levels showed a non-significant reduction at this timepoint, suggesting that longer term expression may be needed for robust DPR reduction in this model (**Fig. 3C**).

**Figure 3.**
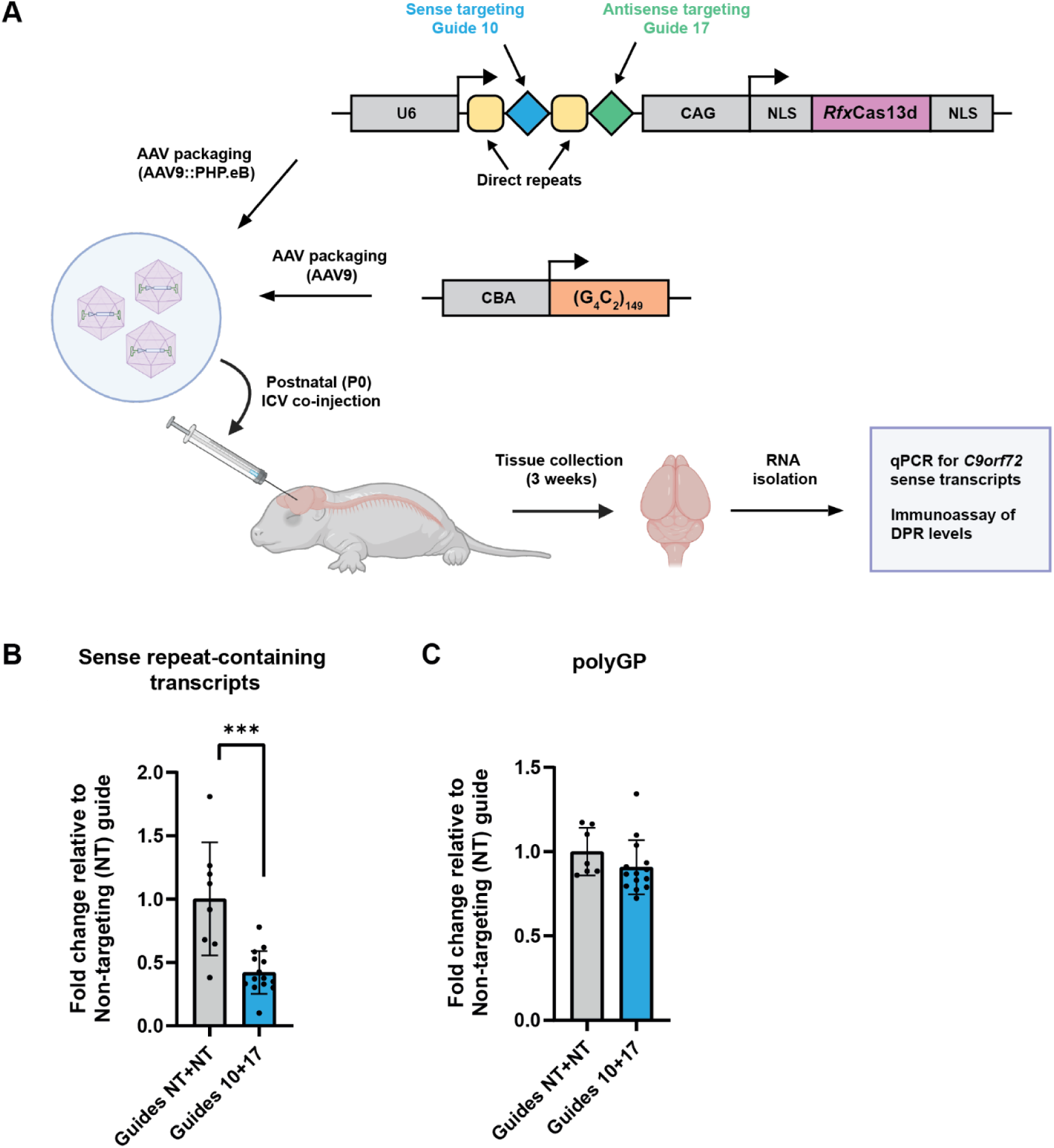
CRISPR-CasRx reduces G_4_C_2_ sense RNA *in vivo* in 149R mice. **(A)** Schematic of experimental paradigm. CasRx PHP.eB AAV and the 149R *C9orf72* AAV9 were co-injected via intracerebroventricular (ICV) injection into P0 C57BL6/J mice. **(B)** RT-qPCR of the sense repeat-containing transcripts from the 149R AAV performed on RNA isolated from the hippocampus 3 weeks post injection. Each data point represents one animal. RT-qPCR data presented as fold change compared to non-targeting (NT+NT) CasRx PHP.eB AAV-treated mice. **(C)** MSD immunoassay of polyGP in bulk hippocampal tissue 3 weeks post-injection. Data given as mean ± S.D, unpaired t-test, ***p<0.001.

### Dual-targeting CRISPR-CasRx effectively reduces sense and antisense repeat RNA in *C9orf72* BAC mice

We next wanted to determine whether our dual-targeting CRISPR-CasRx AAV could reduce both the sense and antisense repeat-containing *C9orf72* transcripts simultaneously *in vivo*. We therefore used a bacterial artificial chromosome (BAC) mouse model that contains the full human *C9orf72* sequence and 500 G_4_C_2_ repeats and thus better emulates the human sense and antisense transcripts ^39^. *C9orf72* BAC transgenic mice were ICV injected at P0 with a CRISPR-CasRx PHP.eB AAV expressing either our sense and antisense targeting gRNAs in an array, or two non-targeting control gRNAs in an array as a control (**Fig. 4A**). To assess overt toxicity due to AAV expression, we measured weights and conducted a general health and welfare assessment ^40,41^ in ICV-injected animals monthly starting at 2 months of age. In either wild-type or BAC transgenics, sense and antisense targeting CasRx AAVs did not significantly alter weight (**Fig. S5A-B**) or induce overt symptoms of sickness (**Fig. S5C-D**). Thus, CasRx AAV delivery was well tolerated. Brain tissue was then harvested at 8.5 months post-injection. *C9orf72* BAC transgenics have been described to occasionally develop promoter methylation and partial transgene silencing ^42^. Therefore, to ensure that any observed change in *C9orf72* RNA or DPR pathology is not due to promoter silencing, we conducted bisulfite pyrosequencing as previously described ^42^ and assessed *C9orf72* promoter DNA methylation levels in the cortices of all animals in the study. No animals in the study had a methylated *C9orf72* promoter (**Fig. 4B**), confirming that any observed changes to RNA and DPR pathologies would be due to CasRx target engagement, rather than transgene silencing. We then assessed bulk cortical tissue for target engagement by RT-qPCR and DPR immunoassay (**Fig. 4C-G**). We observed a significant ∼20% decrease in both sense and antisense repeat-containing transcripts whilst variant 2 was spared (**Fig. 4C-E**). DPR levels were more variable, but were also reduced, with a significant reduction in polyGP (**Fig. 4F-G**). Taken together these data show that our approach can simultaneously reduce both sense and antisense repeat transcripts *in vivo*.

**Figure 4.**
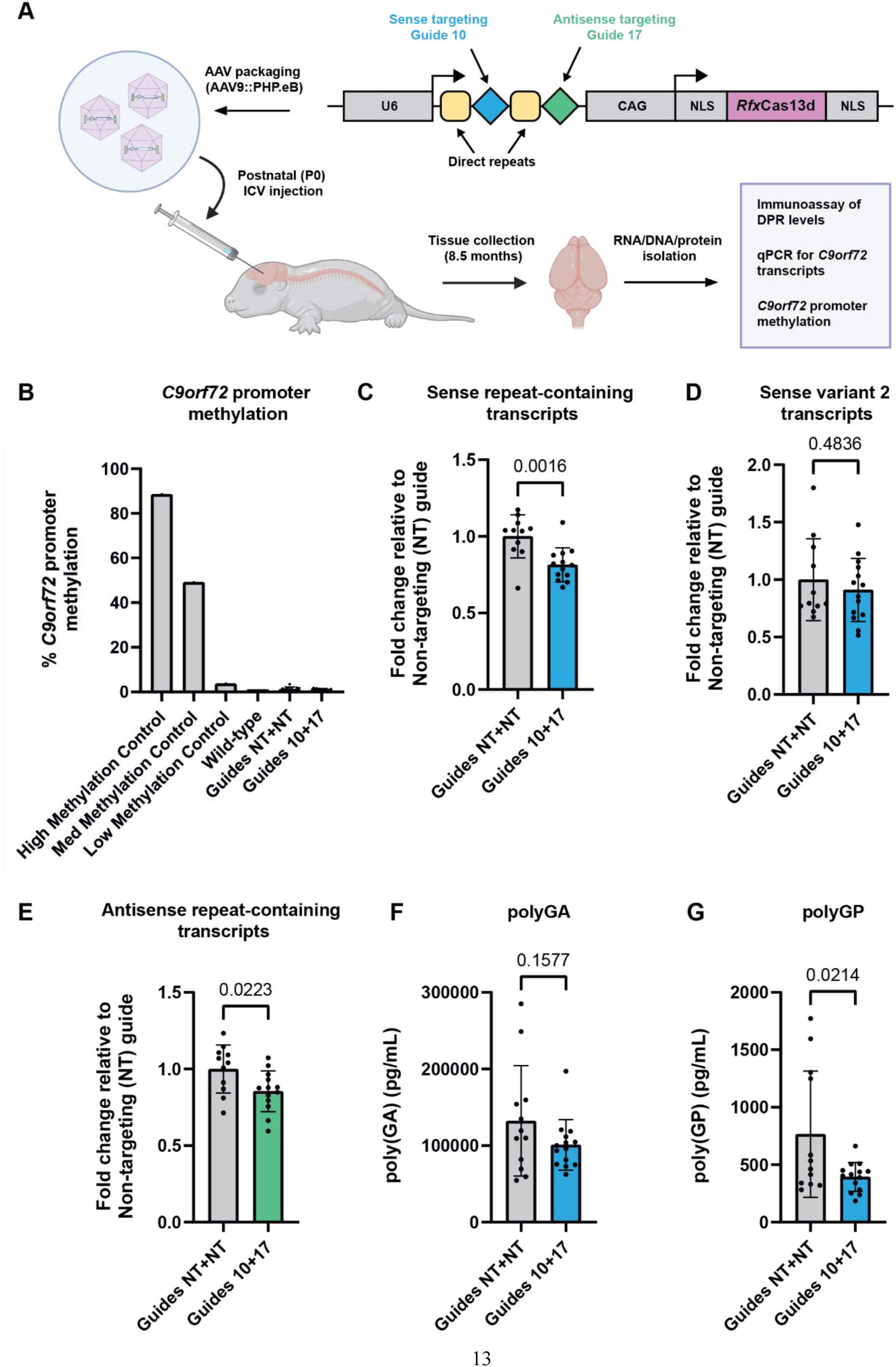
CRISPR-CasRx simultaneously reduces sense and antisense repeat RNAs in *C9orf72* BAC transgenic mice. **(A)** Diagram of experimental paradigm. CasRx PHP.eB AAV (8E+9 vg per animal) was injected via intracerebroventricular (ICV) injection into postnatal day 0 (P0) *C9orf72* BAC mice. Tissue was harvested 8.5 months post-injection. **(B)** *C9orf72* promoter methylation assay of cortical DNA extracted from *C9orf72* BAC mice, demonstrating no overt transgene silencing within the cohort. **(C-E)** RT-qPCRs for *C9orf72* transcripts in *C9orf72* BAC mice treated with CRISPR-CasRx + gRNA AAVs in bulk cortical tissue 8.5 months post-injection. **(C)** RT-qPCR for sense repeat-containing *C9orf72* transcripts (variants 1 and 3). **(D)** RT-qPCR for sense variant 2 *C9orf72* transcripts. **(E)** RT-qPCR for antisense repeat-containing *C9orf72* transcripts. RT-qPCR data presented as fold change compared to non-targeting (NT+NT) gRNA CasRx PHP.eB AAV-treated mice. Data given as mean ± S.D, unpaired t-test. **(E-F)** Levels of **(F)** polyGA and **(G)** polyGP DPRs in bulk cortical tissue from 8.5 month old animals treated with CasRx PHP.eB AAV. Data given as mean ± S.D, n=14 Guide 10+17; n=12 Guide NT+NT, unpaired t-test.

## Discussion

Here we show CasRx can simultaneously target and lower both the sense and antisense *C9orf72* repeat transcripts in transiently expressing cell models, patient iPSC-derived neurons, and *C9orf72* mice without affecting variant 2 expression. We further show that CasRx can mature its own gRNA array and maintain targeting efficiency, allowing for packaging into a single viral vector. A large body of evidence suggests that toxic gain of function is a key driver of *C9orf72*-related ALS/FTD ^4^, however loss or reduction of *C9orf72* has been shown to exacerbate gain of function toxicity ^43,44^. Therefore, an important part of any therapy should be to minimise any further reduction in *C9orf72* expression. To this end, we targeted our guides to the sequence upstream of the repeat expansion in intron 1, thus targeting only those transcripts that contain the repeats. This strategy successfully reduced endogenous repeat-containing transcript variants 1 and 3, whilst sparing variant 2. For any genetic therapy it is vital to minimise off-target effects. CasRx leads to bystander cleavage of RNAs when it is targeted to highly-expressed transcripts, with less bystander cleavage when targeting transcripts with lower expression levels ^34^. We did not observe off-targets using bulk RNA-sequencing when treating *C9orf72* patient iPSC-neurons. This is consistent with the original report of CasRx, which showed no significant off-target transcriptomic changes when targeting *B4GALNT1* in mammalian cells, compared to over 500 off-target changes when targeted using a comparable RNA interference method ^25^. However, more in-depth and sensitive analyses may be required to reveal low-level and unique off-target events.

To date there are no therapies in clinical trial for C9 ALS/FTD that target both the sense and antisense repeat-containing *C9orf72* RNA. This is despite a growing body of evidence suggesting the antisense repeat expansion RNA contributes to disease pathogenesis. This includes the overt toxicity of antisense-derived polyPR in vitro and in vivo ^17–19;^ recent work showing antisense repeat-containing RNA triggers activation of the PKR/eIF2α-dependent integrated stress response independent of DPRs and sense strand-related pathology, leading to aberrant stress granule formation ^21^; and data showing that only ASOs targeting antisense repeats are able to reduce TDP-43 loss of function cryptic splicing changes in *C9orf72* patient iPSC-neurons ^20^. The need to target antisense repeats appears particularly compelling given the recent failures of two clinical trials targeting sense repeats, which did not show clinical efficacy despite reducing sense pathologies (https://investors.biogen.com/news-releases/news-release-details/biogen-and-ionis-announce-topline-phase-1-study-results; https://www.thepharmaletter.com/article/wave-life-sciences-ends-wve-004-program). Our data show that CasRx is one plausible approach for targeting both sense and antisense repeat transcripts.

It has previously been shown that some people already possess antibodies to certain Cas9 orthologs ^45^, although the immune response does not trigger extensive cell damage *in vivo* ^46^. One potential limitation to our current approach is the necessity for long-term expression of CasRx. A recent study identified CasRx-reactive antibodies, and CD4 and CD8 T cell responses in human samples ^47^. This was surprising as CasRx is isolated from *Ruminococcus flavefaciens* strain XPD3002 which is bovine specific and is not present in humans. However CasRx shares ∼35% homology with another Cas13 ortholog isolated from *R. bicirculans* that does colonise humans ^48^. Importantly, there is a precedent that T-cell responses are manageable with immune suppression, suggesting these issues are surmountable ^49^.

Delivery of gene therapies to the CNS remains one of the greatest challenges in the field. In our proof-of-concept experiment in BAC transgenic mice, the decrease in sense and antisense repeat RNA that we observed in bulk cortical tissue was likely limited by the extent of transduction efficiency. The significant reduction observed indicates that there were high levels of knockdown in the cells that were successfully transduced. Thus, delivery with an appropriate capsid and route of administration to enable widespread transduction should provide successful targeting. Our proof-of-concept data suggest CRISPR-CasRx and our gRNAs have potential therapeutic utility via targeting both the sense and antisense repeat-containing *C9orf72* transcripts in a single design small enough to be packaged into an AAV.

## Acknowledgments

We thank Dr Jamie Evans for technical assistance with FACS, Drs Siddharthan Chandran and Bhuvaneish Selvaraj for providing *C9orf72* patient lines, Dr Michael Ward for providing the piggyBac NGN2 constructs and Dr Dieter Edbauer for providing anti-GA monoclonal antibody.

## Funding

Leonard Wolfson Centre for Experimental Neurology, Leonard Wolfson PhD Programme in Neurodegeneration (LK, FK). Neurogenetic Therapies Programme, funded by Sigrid Rausing Trust (AMI, LK). European Research Council, European Union’s Horizon 2020 research and innovation programme grant 648716 – C9ND (AMI). UK Dementia Research Institute, funded by UK Medical Research Council, Alzheimer’s Society, and Alzheimer’s Research UK (AMI). EMBO Postdoctoral Fellowship (AJC). Live Like Lou Foundation Postdoctoral Fellowship (AJC). Wolfson-Eisai Neurodegeneration University College London PhD programme, funded by Eisai (RC). Alzheimer’s UK grant ARUK-DC2021-039 (TMR, AM). Wellcome Investigator Award 217150/Z/19/Z (JR). UK Medical Research Council MR/N026101/1, MR/T044853/1, MR/X004724/1, MR/R025134/1, MR/R015325/1, MR/S009434/1 (AAR). The Wellcome Trust Institutional Strategic Support Fund/UCL Therapeutic Acceleration Support (TAS) Fund 204841/Z/16/Z (AAR). The Sigrid Rausing Trust and the Jameel Education Foundation (AAR). NIHR Great Ormond Street Hospital Biomedical Research Centre 562868 (AAR).

## Author contributions

Conceptualization: LK, AMI. Methodology: LK, DV, AJC, AMI, BVH, AM, TMR, LP, MW, JR, AAR. Investigation: LK, DV, AJC, MC, PS, BM, TGM, LY, FK, BVH, RC, ANC. Analysis:

LK, DV, AJC, MC, FK, JR. Visualization: LK, DV, BH, FK. Writing – original draft: LK, AMI, DV, AJC. Writing – review & editing: all authors.

## Competing interests

LK and AMI are co-inventors listed on UK Patent Application No. 2105455.6, International application No. PCT/EP2022/060296 which are related to the presented data. All other authors declare they have no competing interests.

## Data and materials availability

All data are available in the main text or the supplementary materials or are available on request.

## Materials and methods

### Ethics statement

*C9orf72* patient iPSC lines were kindly provided by Prof Siddharthan Chandran and Dr Bhuvaneish Selvaraj, University of Edinburgh and have been previously described ^50^; they were collected with prior informed patient consent and derived from biopsied fibroblasts. Ethical approval was received from the NHS Health Research Authority East of England - Essex Research Ethics Committee (REC reference 18/EE/0293). Animals were maintained and experimental procedures performed in accordance with the UK Animal Scientific Procedure Act 1986, under project and personal licenses issued by the UK Home Office.

### Plasmid Construction

#### NanoLuc Reporter Plasmids

The sense NLuc reporter was as previously published ^31^. To generate an antisense RAN translation NLuc reporter construct, 680 nucleotides of the endogenous *C9orf72* sequence upstream of the antisense repeats was synthesised and cloned into the NLuc backbone expression vector and the G_4_C_2_ repeats from the sense NLuc plasmid were then cloned in the reverse direction using NotI and BspQI to make C_4_G_2_ repeats with the endogenous upstream sequence.

A repeat insert of ∼55 C_4_G_2_ repeats was confirmed by sizing on an agarose gel.

#### gRNA Plasmids

Guides were designed using BLAST to identify off-targets and secondary structure scores were predicted using RNAfold Webserver (University of Vienna). We used a previously published non-targeting control guide ^24^. Annealed guide oligonucleotides were designed to have overhangs for cloning into their respective gRNA expression backbone vectors using BbsI.

#### gRNA-CasRx Lentiviral Plasmids

The CasRx lentiviral vector (pXR001) was used as a backbone and primers were designed to PCR out the U6 promoter, direct repeats, and gRNA from the guide expressing vector (pXR003) with PacI restriction site overhangs to allow for cloning into the CasRx backbone and confirmed by sequencing.

#### CasRx AAV-PHP.eB Plasmids

An insert containing our CRISPR pre-gRNA array and CasRx driven by a 643 bp CAG variant promoter ^51^ was synthesised (GeneArt, Thermo Fisher) and was cloned into an AAV backbone vector using restriction sites NotI and AscI. Golden Gate cloning utilising type IIs restriction enzyme BsmBI was then used to clone in our guide array of choice (guides 10 + 17). Correct insertion of the CasRx insert and gRNA array as well integrity of the ITRs was confirmed via Sanger sequencing.

### HEK293T cell culture

HEK293T cells (a kind gift from the UCL Drug Discovery Institute) were maintained in DMEM media supplemented with 10% FBS, 4.5 g/L glucose, 110 mg/L sodium pyruvate, 1x GlutaMAX and kept at 37°C with 5% CO2 to ensure physiological temperature and pH. Cells were maintained up to a confluency of 90% and then dissociated and passaged with 0.05% Trypsin-EDTA.

### Lentivirus and AAV production

Low passage HEK293T cells were cultured as described above and plated in T175 flasks at 50% confluency 24 hours prior to transfection with 14.1 µg of PAX lentiviral packaging vector, 9.36 µg of VSV.G lentiviral enveloping vector and 14.1 µg of lentivirus plasmid of interest using Lipofectamine 3000. 24 hours after transfection, cell media was replaced with NGN2 neuronal induction media. The media was collected 24 hours later and filtered to remove cell debris via a 45 µm filter before being aliquoted and stored at −80°C until needed.

To produce PHP.eB serotype AAVs containing our CasRx and gRNA inserts, low passage HEK293T cells were triple transfected with plasmids containing our CasRx expression cassette flanked by 5’ and 3’ inverted terminal repeats (ITRs), a PHP.eB Cap plasmid, and a helper plasmid in a 1:1:3 plasmid copy number ratio. 72 hours post-transfection supernatant and cells were collected and virus was purified via ultracentrifugation in an increasing gradient of iodixanol at 200,000 x g for 3 hours with isolation of viral particles in the 40% fraction. This fraction was then filtered, concentrated, and titered via Vivaspin^TM^ 20 concentrator and gel electrophoresis. PHP.eB AAVs were made at the GeneTxNeuro Vector Core Facility at the UCL School of Pharmacy and a second batch was outsourced to the viral vector facility at ETH Zurich. 149R AAV9 viral particles were generated as previously described ^38^.

### NanoLuciferase reporter assays

For luciferase assays, HEK293T were plated at a density of 30,000 cells per well (96 well plate) and transiently transfected with 100 ng of Cas13 gRNA plasmids, 25 ng of Cas13 plasmid, 12.5 ng of Firefly luciferase expression plasmid, and 2.5 ng of RAN translation sense or antisense NanoLuciferase reporter plasmids (S92R-NL and AS55R-NL respectively). Transfection reagents were added directly to the media and left on for the duration of the experiment. Each experiment consists of 3-5 technical replicate wells per condition. 48 hours post-transfection both Firefly and NanoLuciferase signals were measured using the Nano-Glo Dual Luciferase Assay (Promega) according to manufacturer’s instructions, on the FLUOstar Omega (BMG Labtech) with a threshold of 80% and a gain of 2000 for both readings. The NanoLuciferase reading was normalised to the Firefly luciferase reading for each well to control for variable transfection efficiencies.

### Combined single molecule RNA FISH and immunocytochemistry

For RNA fluorescent *in situ* hybridisation (RNA FISH) in HEK293T cells, cells were plated at 25,000 cells per well in a 96 well plate and transfected with 100 ng of CasRx gRNA plasmids, 25 ng of CasRx plasmids, 12.5 ng of Firefly luciferase expression plasmid, and 2.5 ng of RAN translation sense or antisense NanoLuciferase reporter plasmids. Each plate contained 3-5 technical replicates per condition. Cells were fixed 48 hours post-transfection for 7 minutes using 4 % Paraformaldehyde (PFA) with 10% methanol, diluted in PBS. Cells were then dehydrated via 70% and 100% ethanol washes and frozen at −80°C in 100% ethanol until needed. Cells were rehydrated with 70% ethanol and washed for 5 minutes at room temperature in pre-hybridisation solution (40% formamide, 2× SSC, 10% dextran sulphate, 2 mM vanadyl ribonucleoside complex). Cells were permeabilised with 0.2% Triton X-100 for 10 minutes. Cells were incubated at 60°C in pre-hybridisation solution for 45 minutes. Locked nucleic acid (LNA) probes (Qiagen) to detect either sense (5’ TYE563-labelled CCCCGGCCCCGGCCCC) or antisense (5’ TYE563-labelled GGGGCCGGGGCCGGGG) RNA-foci were then added to the pre-hybridisation solution at 40 nM and cells were kept in the dark at 60°C or 66°C (for sense or antisense probes, respectively) for 3 hours or overnight. Cells were then washed with 0.2% Triton-X100 in 2x SSC for 5 minutes at room temperature followed by 30 minutes at 60°C. One additional wash in 0.2x SSC at 60°C for 30 minutes was performed prior to application of 647-conjugated HA antibody (in 0.2x SSC with 1% BSA at 1:1000) for detection of HA-tagged CasRx. This was incubated overnight at 4°C and then washed with 0.2x SSC for 20 minutes at room temperature. Hoechst was added at 1:5000 in 0.2x SSC for 10 minutes to detect cell nuclei. Hoechst solution was removed, and cells were left in 0.2x SSC at 4°C protected from light until imaging.

### iPSC culture

The three *C9orf72* patient-derived iPSC lines used are described in **Sup. Table 1**. iPSCs were cultured on Geltrex-coated wells, fed with fresh E8 media (Gibco) daily, and passaged with EDTA (0.5 mM).

### Generation of NGN2-inducible iPSC neurons

*C9orf72* patient-derived iPSCs expressing doxycycline-inducible NGN2 iPSCs were generated by piggyBac integration as previously described ^32,33^. Cells underwent FACS using the BFP in the NGN2 construct to ensure a pure population of NGN2 iPSCs.

### Differentiation of NGN2-inducible iPSCs into cortical-like neurons

iPSCs were differentiated to cortical-like neurons via dox-inducible NGN2 expression as previously described ^52^. Briefly, the cells were cultured to ∼80% confluency prior to differentiation. On DIV0 of neuronal differentiation, NGN2 iPSCs were dissociated and lifted with Accutase (Gibco). Cells were replated at desired ratio (3.75 x 10^5^ cells per well/of a 6-well plate) on Geltrex-coated wells in neuronal induction media: DMEM-F12 (Gibco) containing 1x N2 (Thermo Fisher Scientific), 1x Glutamine (Gibco), 1x HEPES (Gibco), 1x NEAA (Gibco), doxycycline (2µg/mL) and 10 µM Rock inhibitor Y-27632 (DIV0 only; Tocris). Cells were washed with PBS the following day to remove cell debris and fresh induction media was added without Rock inhibitor. On DIV3, induction media was removed and replaced with neuronal maintenance media: Neurobasal (Gibco), supplemented with 1x B27 (Gibco), 10 ng/mL BDNF (PeproTech), 10 ng/mL NT-3 (PreproTech) and 1 µg/mL laminin. Cells were given a half-media change on DIV4 with fresh maintenance media and then used for downstream processing on DIV5 as indicated.

### Lentiviral Delivery of CasRx and guide RNA to NGN2 iPSC-neuron and fluorescence-activated cell sorting

NGN2 iPSCs were treated with lentiviruses on DIV0 of the differentiation protocol, 2 hours post dissociation with Accutase. 24 hours later, media was replaced with fresh induction media and the differentiation protocol followed. At DIV5, cells were lifted and dissociated with Accutase for downstream analyses. For FACS, cells were pelleted via centrifugation prior to resuspension in FACS buffer (PBS + 1% BSA + 2mM EDTA). Cell clumps were removed via a 100 µm cell strainer and cells were resuspended in FACS buffer at 3 x 10^6^ cells/mL. Cells were sorted at the UCL Flow Cytometry Core Facility using the BD FACSAria^TM^ III Cell Sorter. Sorted cells were collected and pelleted via centrifugation and snap frozen for use in downstream analyses.

### Protein extraction and polyGA and polyGP Meso Scale Discovery (MSD) immunoassays

To extract protein from the NGN2-iPSC neurons for immunoassays, FACS purified cell pellets were incubated in lysis buffer (2% SDS + protease inhibitors in RIPA buffer) for 5 minutes at room temperature prior to sonication for 3 x 5 seconds at 30 amp at 4°C. Samples then underwent centrifugation at 17,000 x g at 4°C for 20 minutes to remove cell debris. To extract protein from *C9orf72* mouse tissue, 100 mg of frozen brain was taken per mouse sample and lysed with 0.9 x tissue mass of lysis buffer (1x RIPA buffer with 2% SDS and 2x protease inhibitor cocktail) and was homogenised using a TissueRuptor II (Qiagen). Zebrafish larvae were similarly extracted, with 0.9 x tissue mass of lysis buffer (1x RIPA buffer with 2% SDS and 2x protease inhibitor cocktail) and TissueRuptor II (Qiagen) homogenization. Samples were then sonicated at 4°C for 3 x 20 seconds at 30% amplitude. Samples were centrifuged at 20,000 x g for 20 minutes at 4°C and supernatant was collected. Protein concentration was determined via BCA assay (Thermo Fisher) according to manufacturer’s instructions. MSD assays were performed as previously described ^53,54^. For polyGP, capture and detection antibody was the same affinity-purified custom anti-(GP)8 antibody (Eurogentec, biotinylated for detector). For polyGA, capture antibody was anti-GA (Millipore, # 5E9-3226534) and detector antibody was monoclonal anti-GA 5F2 kindly gifted by Prof Dieter Edbauer (DZNE Munich) and biotinylated in-house.

### RNA extraction and RT-qPCR

Total RNA was extracted from FACS purified iPSC-neurons and frozen mouse brain tissue using the ReliaPrep RNA cell kit and ReliaPrep RNA tissue extraction kit (Promega) respectively.

RNA samples underwent reverse transcription using SuperScript^TM^ IV VILO^TM^ Master Mix (ThermoFisher) with random hexamers to produce cDNA according to manufacturer’s instructions. qPCR analysis was performed using SYBR Green Master Mix (ThermoFisher) for all qPCRs except *C9orf72* variant 2 expression levels for which TaqMan probes were used (sequences provided in Table S2) and measured with a LightCycler^TM^ (Roche). qPCR was performed in technical triplicate with data normalised to GAPDH expression levels.

### RNA sequencing and analysis

RNA library was prepared for sequencing with KAPA RNA HyperPrep with RiboErase. RNA-sequencing (NextSeq 2000) was carried out at a depth of 20 million reads per sample. RNA-seq reads were pseudoaligned to the reference human transcriptome (Ensembl GRCh38 v96) using kallisto (v.0.46.0) ^55^. Differential gene expression analysis was performed using the DEseq2 package (v1.36.0) ^56^ by importing the estimated abundance data using the tximport package ^57^. The transcripts of *C9orf72* were categorised into two groups based on the presence or absence of intron 1. The threshold for differentially expressed genes was an adjusted p-value of less than 0.05 and log2FoldChange>1. R packages EnhancedVolcano (v.1.14.0), ggplot2 (v.3.4.1), and ggsci (v.2.9) were used for data visualisation.

### Cell viability assay

NGN2 iPSCs were plated on Geltrex-coated 96-well microplates and differentiation and transduction protocols were followed as above. Cell viability was assessed 5 days post-transduction via CellTiter-Fluor^TM^ assay according to manufacturer’s instructions. 20 mg/mL digitonin (Merck) was added as a positive control. Fluorescent signals (380-400 nm_Ex_ / 505 nm_Em_) were determined via analysis with the PHERAstar^TM^ microplate reader (BMG Labtech).

### Zebrafish experiments

Adult zebrafish were reared by University College London’s Fish Facility on a 14 hr:10 hr light:dark cycle. All experiments used the progeny of an outcross of heterozygous *Tg(ubi:G_4_C_2_×45; hsp70:DsRed)* ^35^ to wild type (AB × Tup LF). To obtain eggs, pairs of one female and one male were isolated in breeding boxes overnight, separated by a divider. Around 9 AM (lights on) the next day, the dividers were removed, and eggs were collected 7–10 min later. The embryos were then raised in 10-cm Petri dishes filled with fish water (0.3 g/L Instant Ocean) in a 28.5°C; incubator on a 14 hr:10 hr light:dark cycle. Debris and dead or dysmorphic embryos were removed every other day with a Pasteur pipette under a bright-field microscope and the fish water replaced. At the end of the experiments, larvae were euthanised with an overdose of 2-phenoxyethanol (ACROS Organics). Experimental procedures were in accordance with the Animals (Scientific Procedures) Act 1986 under Home Office project licences PA8D4D0E5 and PP6325955 awarded to Jason Rihel. Adult zebrafish were kept according to FELASA guidelines_58._

In the experiment in **Fig. S4C,D**, each clutch was from a unique pair of parents. In each CasRx experiment (**FIG S4E–I**), single-cell embryos from different pairs were pooled and injected in the yolk with 10 pg of CasRx targeting or non-targeting plasmid and 500 pg of Tol2 mRNA in 1 nL. Only larvae with strong GFP expression were kept for the behaviour tracking experiment. At 5 dpf, individual larvae were transferred to the wells of clear 96-square well plates (Whatman). To avoid any potential localisation bias during the tracking, wild-type and *ubi:G_4_C_2_×45* heterozygous larvae (identified by DsRed expression) were plated in alternating columns of the 96-well plate. The plates were placed in ZebraBoxes (Viewpoint Behavior Technology). From each well, the video-tracking software (ZebraLab, Viewpoint Behavior Technology) recorded the number of pixels that changed intensity between successive frames. To be counted, a pixel must have changed grey value above a sensitivity threshold, which was set at 20. The metric, termed Δ pixel, describes each animal’s behaviour over time as a sequence of zeros and positive values, denoting if the larva was still or moving. Tracking was performed at 25 frames per second on a 14 hr:10 hr light:dark cycle for ∼65 hours, generating sequences of roughly 5,850,000 Δ pixel values per animal. In plots (**Fig. S4C,H**), the first day and night (5 dpf) are removed as acclimatisation period. The day light level was calibrated at 555 lux with a RS PRO RS-92 light meter (RS Components). Night was in complete darkness with infrared lighting for video recording. Both mornings shortly after 9 AM (lights on), the wells were manually topped-up with fish water to counteract water evaporation. At the end of the tracking, any larva that did not appear healthy was excluded from subsequent analysis. Larvae were anaesthetised then snap-frozen in liquid nitrogen for DPR measurements.

Behavioural data analysis was performed using the FramebyFrame R package v0.13.0 ^37^. In **Fig. S4D,I**, time spent active per day was statistically compared between genotypes using linear mixed effects (LME) modelling implemented in the lmer function of the R package lme4 v1.1.31^59^. The command to create the LME model was:

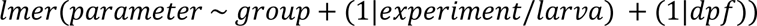

Each group was then compared to the reference group using estimated marginal means implemented in the R package emmeans, which provided the p-value.

For the pictures in **Fig. S4A,F**, larvae were anaesthetised and mounted in 1% low melting point agarose (Sigma-Aldrich) in fish water. Pictures were then taken with an Olympus MVX10 microscope connected to a computer with the software cellSens (Olympus).

### *C9orf72* 149R and BAC mouse model housing, AAV administration, and tissue harvesting

C57BL6/J mice were used for 149R experiments. Bacterial artificial chromosome (BAC) *C9orf72* mice ^39^ in the FVB/NJ background were obtained from the Jackson Laboratory (RRID:IMSR_JAX:029099) and were PCR genotyped using the Jackson Laboratory protocol. Within 24 hours of birth, pups were manually injected with AAVs via intracerebroventricular (ICV) injection into both hemispheres. Briefly, AAVs were diluted in sterile PBS to a final volume of 5 µL per animal. P0 pups were anaesthetized with isoflurane, after which a calibrated Hamilton 10 µL syringe was inserted into the skull approximately 2/5 of the distance from lambda to the eye and at a depth of approximately 2 mm. 2.5 µL of AAV/PBS solution was injected slowly into each hemisphere. After injection, pups were allowed to recover on a heat pad and then returned to the dam. 149R AAV was administered at 6E+10 vg per animal as described previously ^38^, CasRx PHP.eB AAVs were administered at 8E+9 vg per animal. For tissue collection, animals were anaesthetised with isoflurane and perfused with ice-cold PBS. Brains were immediately collected, dissected, snap-frozen on dry ice, and stored at −80°C until use. All *in vivo* experiments used both male and female animals in Mendelian ratios.

### *C9orf72* BAC promoter methylation assay

Promoter methylation for BAC transgenics was assessed via bisulfite pyrosequencing, conducted by EpigenDx (Assay ADS3232-FS1) as previously described ^42^. ADS3232-FS1 covers 16 CpG’s within the *C9orf72* promoter, spanning the transcription start site, from base pairs −55 to +125.

### Mouse clinical scoring

To assess general health and welfare of ICV-injected animals post-AAV delivery, we conducted clinical scoring monthly beginning at 2 months of age until the end of the study (8.5 months of age). The composite clinical score (18-point scale) is adapted from similar health assessments ^40,41,60^ and assesses 6 aspects of general health and fitness: weight (% change from previous timepoint), posture, coat/grooming, eye squinting, activity, and breathing. Each component is scored 0-3 and are added together for a composite score of 0-18. A composite score of 0-3 is considered healthy/no burden of distress, while 4-8 is mild, 9-12 is moderate, and 13-18 is severe requiring culling.

### Imaging techniques and quantification

RNA-FISH and immunocytochemistry experiments were all imaged using the automated Opera Phenix ^TM^ high-throughput confocal imaging platform (Perkin Elmer). Dual RNA-FISH and immunocytochemistry images were analysed using Columbus 2.8 (PerkinElmer): RNA FISH load was determined by calculating the integrated intensity of nuclear RNA puncta by multiplying spot intensity by total spot load per CasRx positive cell (as determined by nuclear HA positivity). Transfection efficiency images were taken using the IncuCyte S3 live cell imager (Essen BioScience). Immunohistochemistry images of mouse sections were obtained using the AxioScan^TM^ (Zeiss).

### Quantification and statistical analysis

Biologically independent experiment replicates (N number) are indicated in figure legends. Within each biological repeat, there was minimum of 3 technical replicates. Technical replicates are displayed on graphs as well biological replicates. Statistical analyses were only performed on experiments with 3 biologically independent replicates. All datasets conformed to Gaussian distribution as determined by Shapiro-Wilk normality testing, therefore parametric one-way or two-way ANOVA tests were carried out with Holm-Sidak post-hoc analysis (*p*=0.05) to determine statistical significance between test groups. Student’s T-test was used to determine significance between two test groups (*p*=0.05). When statistical significance is indicated in figures, \**p*<0.05; \*\**p*<0.01; \*\*\**p*<0.001; \*\*\*\**p*<0.0001. All statistical tests were carried out using GraphPad Prism 8.4.1. All data in text and figures, unless otherwise stated, are given as fold-change compared to non-targeting guide control. Data displayed as mean values ± standard deviation (S.D.). Workflow diagrams made with permissioned use from Biorender.com.

---

**Supplementary Table 1.** iPSC donor information.

| **iPSC line nomenclature** | **Gender (M/F)** | **Clinical diagnosis** | **Age of onset (yrs)** | **Age of death (yrs)** | **No. of G<sub>4</sub>C<sub>2</sub> repeats** | **Source of iPSC cells** |
| --- | --- | --- | --- | --- | --- | --- |
| C9 Line 1 | F | ALS/FTD | NA | NA | ~750 | <sup>50</sup> |
| C9 Line 2 | M | ALS | 52 | 58 | ~640 | <sup>50</sup> |
| C9 Line 3 | M | ALS | NA | NA | ~960 | <sup>50</sup> |

**Supplementary Table 2.**
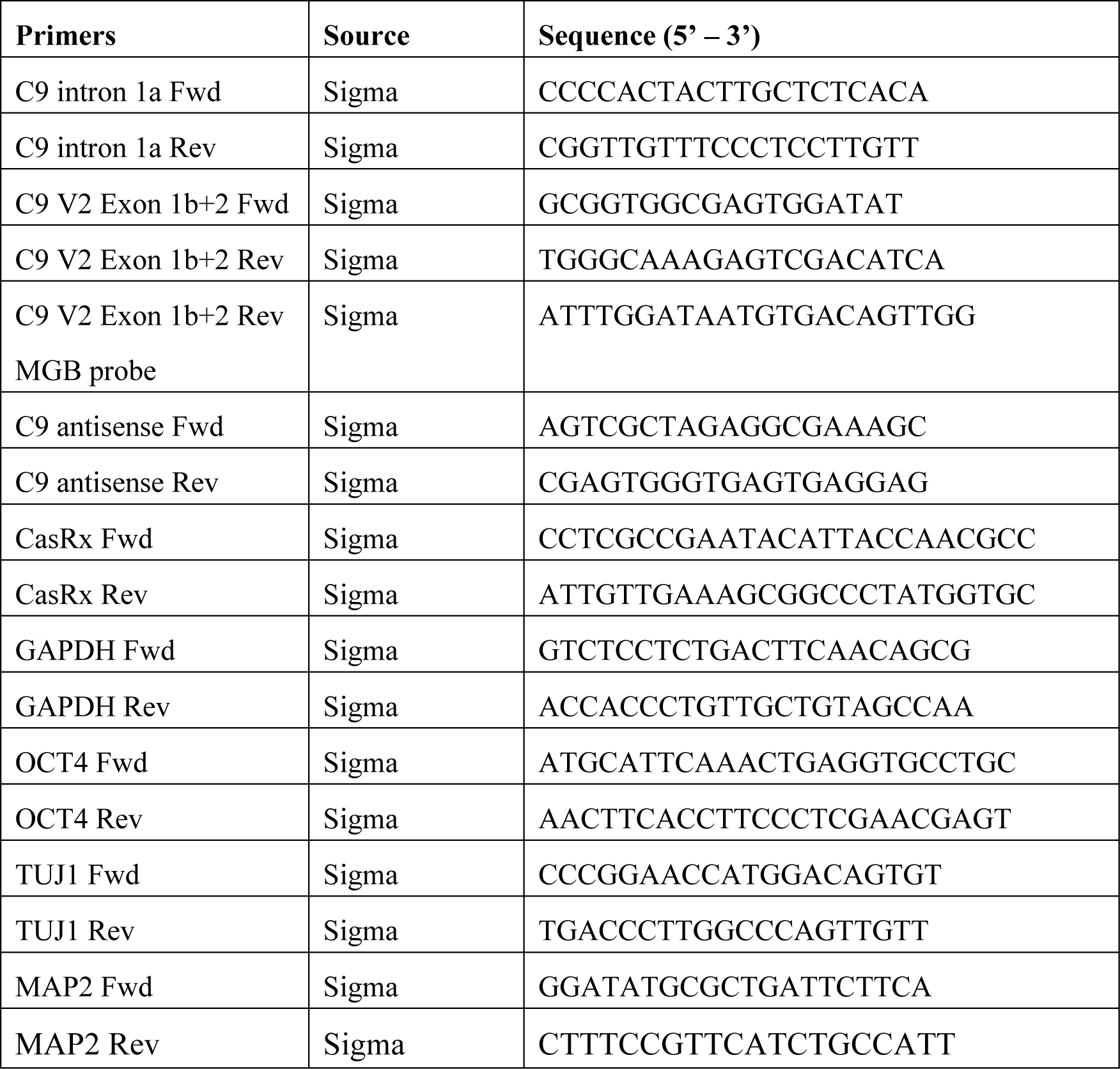
List of qPCR primers.

## Notes

### Competing Interest Statement

Liam Kempthorne and Adrian Isaacs are co-inventors listed on UK Patent Application No. 2105455.6, International application No. PCT/EP2022/060296 which are related to the presented data. All other authors declare they have no competing interests.

## References

1. Burrell, J.R., Halliday, G.M., Kril, J.J., Ittner, L.M., Götz, J., Kiernan, M.C., and Hodges, J.R. (2016). The frontotemporal dementia-motor neuron disease continuum. Lancet (London, England) 388, 919–931. 10.1016/S0140-6736(16)00737-6.

2. DeJesus-Hernandez, M., Mackenzie, I.R., Boeve, B.F., Boxer, A.L., Baker, M., Rutherford, N.J., Nicholson, A.M., Finch, N.A., Flynn, H., Adamson, J., et al. (2011). Expanded GGGGCC Hexanucleotide Repeat in Noncoding Region of C9ORF72 Causes Chromosome 9p-Linked FTD and ALS. Neuron 72, 245–256. 10.1016/j.neuron.2011.09.011.

3. Renton, A.E., Majounie, E., Waite, A., Simón-Sánchez, J., Rollinson, S., Gibbs, J.R., Schymick, J.C., Laaksovirta, H., van Swieten, J.C., Myllykangas, L., et al. (2011). A Hexanucleotide Repeat Expansion in C9ORF72 Is the Cause of Chromosome 9p21-Linked ALS-FTD. Neuron 72, 257–268. 10.1016/j.neuron.2011.09.010.

4. Balendra, R., and Isaacs, A.M. (2018). C9orf72-mediated ALS and FTD: multiple pathways to disease. Nat. Rev. Neurol. 14, 544–558. 10.1038/s41582-018-0047-2.

5. Mizielinska, S., Lashley, T., Norona, F.E., Clayton, E.L., Ridler, C.E., Fratta, P., and Isaacs, A.M. (2013). C9orf72 frontotemporal lobar degeneration is characterised by frequent neuronal sense and antisense RNA foci. Acta Neuropathol. 126, 845–857. 10.1007/s00401-013-1200-z.

6. Zu, T., Liu, Y., Bañez-Coronel, M., Reid, T., Pletnikova, O., Lewis, J., Miller, T.M., Harms, M.B., Falchook, A.E., Subramony, S.H., et al. (2013). RAN proteins and RNA foci from antisense transcripts in C9ORF72 ALS and frontotemporal dementia. Proc. Natl. Acad. Sci. U. S. A. 10.1073/pnas.1315438110.

7. Gendron, T.F., Bieniek, K.F., Zhang, Y.J., Jansen-West, K., Ash, P.E.A., Caulfield, T., Daughrity, L., Dunmore, J.H., Castanedes-Casey, M., Chew, J., et al. (2013). Antisense transcripts of the expanded C9ORF72 hexanucleotide repeat form nuclear RNA foci and undergo repeat-associated non-ATG translation in c9FTD/ALS. Acta Neuropathol. 10.1007/s00401-013-1192-8.

8. Lagier-Tourenne, C., Baughn, M., Rigo, F., Sun, S., Liu, P., Li, H.R., Jiang, J., Watt, A.T., Chun, S., Katz, M., et al. (2013). Targeted degradation of sense and antisense C9orf72 RNA foci as therapy for ALS and frontotemporal degeneration. Proc. Natl. Acad. Sci. U. S. A. 10.1073/pnas.1318835110.

9. Ash, P.E.A., Bieniek, K.F., Gendron, T.F., Caulfield, T., Lin, W.L., DeJesus-Hernandez, M., Van Blitterswijk, M.M., Jansen-West, K., Paul, J.W., Rademakers, R., et al. (2013). Unconventional translation of C9ORF72 GGGGCC expansion generates insoluble polypeptides specific to c9FTD/ALS. Neuron 77, 639–646. 10.1016/J.NEURON.2013.02.004.

10. Mori, K., Weng, S.M., Arzberger, T., May, S., Rentzsch, K., Kremmer, E., Schmid, B., Kretzschmar, H.A., Cruts, M., Van Broeckhoven, C., et al. (2013). The C9orf72 GGGGCC repeat is translated into aggregating dipeptide-repeat proteins in FTLD/ALS. Science 339, 1335–1338. 10.1126/SCIENCE.1232927.

11. Mori, K., Arzberger, T., Grässer, F.A., Gijselinck, I., May, S., Rentzsch, K., Weng, S.M., Schludi, M.H., Van Der Zee, J., Cruts, M., et al. (2013). Bidirectional transcripts of the expanded C9orf72 hexanucleotide repeat are translated into aggregating dipeptide repeat proteins. Acta Neuropathol. 126, 881–893. 10.1007/S00401-013-1189-3.

12. Bush, J.A., Aikawa, H., Fuerst, R., Li, Y., Ursu, A., Meyer, S.M., Benhamou, R.I., Chen, J.L., Khan, T., Wagner-Griffin, S., et al. (2021). Ribonuclease recruitment using a small molecule reduced c9ALS/FTD r(G4C2) repeat expansion in vitro and in vivo ALS models. Sci. Transl. Med. 13. 10.1126/SCITRANSLMED.ABD5991.

13. Jiang, J., Zhu, Q., Gendron, T.F., Saberi, S., McAlonis-Downes, M., Seelman, A., Stauffer, J.E., Jafar-nejad, P., Drenner, K., Schulte, D., et al. (2016). Gain of Toxicity from ALS/FTD-Linked Repeat Expansions in C9ORF72 Is Alleviated by Antisense Oligonucleotides Targeting GGGGCC-Containing RNAs. Neuron. 10.1016/j.neuron.2016.04.006.

14. Liu, Y., Dodart, J.-C., Tran, H., Berkovitch, S., Braun, M., Byrne, M., Durbin, A.F., Hu, X.S., Iwamoto, N., Jang, H.G., et al. (2021). Variant-selective stereopure oligonucleotides protect against pathologies associated with C9orf72-repeat expansion in preclinical models. Nat. Commun. 12. 10.1038/s41467-021-21112-8.

15. Martier, R., Liefhebber, J.M., Miniarikova, J., van der Zon, T., Snapper, J., Kolder, I., Petry, H., van Deventer, S.J., Evers, M.M., and Konstantinova, P. (2019). Artificial MicroRNAs Targeting C9orf72 Can Reduce Accumulation of Intra-nuclear Transcripts in ALS and FTD Patients. Mol. Ther. - Nucleic Acids. 10.1016/j.omtn.2019.01.010.

16. Cabrera, G.T., Meijboom, K.E., Abdallah, A., Tran, H., Foster, Z., Weiss, A., Wightman, N., Stock, R., Gendron, T., Gruntman, A., et al. (2023). Artificial microRNA suppresses C9ORF72 variants and decreases toxic dipeptide repeat proteins in vivo. Gene Ther. 10.1038/S41434-023-00418-W.

17. Mizielinska, S., Grönke, S., Niccoli, T., Ridler, C.E., Clayton, E.L., Devoy, A., Moens, T., Norona, F.E., Woollacott, I.O.C., Pietrzyk, J., et al. (2014). C9orf72 repeat expansions cause neurodegeneration in Drosophila through arginine-rich proteins. Science 345, 1192– 1194. 10.1126/science.1256800.

18. Wen, X., Tan, W., Westergard, T., Krishnamurthy, K., Markandaiah, S.S., Shi, Y., Lin, S., Shneider, N.A., Monaghan, J., Pandey, U.B., et al. (2014). Antisense proline-arginine RAN dipeptides linked to C9ORF72-ALS/FTD form toxic nuclear aggregates that initiate in vitro and in vivo neuronal death. Neuron 84, 1213–1225. 10.1016/J.NEURON.2014.12.010.

19. Zhang, Y.J., Guo, L., Gonzales, P.K., Gendron, T.F., Wu, Y., Jansen-West, K., O’Raw, A.D., Pickles, S.R., Prudencio, M., Carlomagno, Y., et al. (2019). Heterochromatin anomalies and double-stranded RNA accumulation underlie C9orf72 poly(PR) toxicity. Science 363. 10.1126/SCIENCE.AAV2606.

20. Rothstein, J.D., Baskerville, V., Rapuri, S., Mehlhop, E., Jafar-Nejad, P., Rigo, F., Bennett, F., Mizielinska, S., Isaacs, A., and Coyne, A.N. (2024). G2C4 targeting antisense oligonucleotides potently mitigate TDP-43 dysfunction in human C9orf72 ALS/FTD induced pluripotent stem cell derived neurons. Acta Neuropathol. 147. 10.1007/s00401-023-02652-3.

21. Parameswaran, J., Zhang, N., Braems, E., Tilahun, K., Pant, D.C., Yin, K., Asress, S., Heeren, K., Banerjee, A., Davis, E., et al. (2023). Antisense, but not sense, repeat expanded RNAs activate PKR/eIF2α-dependent ISR in C9ORF72 FTD/ALS. Elife 12. 10.7554/eLife.85902.

22. East-Seletsky, A., O’Connell, M.R., Knight, S.C., Burstein, D., Cate, J.H.D., Tjian, R., and Doudna, J.A. (2016). Two distinct RNase activities of CRISPR-C2c2 enable guide-RNA processing and RNA detection. Nature 538, 270–273. 10.1038/NATURE19802.

23. Abudayyeh, O.O., Gootenberg, J.S., Essletzbichler, P., Han, S., Joung, J., Belanto, J.J., Verdine, V., Cox, D.B.T., Kellner, M.J., Regev, A., et al. (2017). RNA targeting with CRISPR-Cas13. Nature 550, 280–284. 10.1038/nature24049.

24. Cox, D.B.T., Gootenberg, J.S., Abudayyeh, O.O., Franklin, B., Kellner, M.J., Joung, J., and Zhang, F. (2017). RNA editing with CRISPR-Cas13. Science (80-.). 358, 1019–1027. 10.1126/science.aaq0180.

25. Konermann, S., Lotfy, P., Brideau, N.J., Oki, J., Shokhirev, M.N., and Hsu, P.D. (2018). Transcriptome Engineering with RNA-Targeting Type VI-D CRISPR Effectors. Cell 173, 665–676.e14. 10.1016/j.cell.2018.02.033.

26. Yan, W.X., Chong, S., Zhang, H., Makarova, K.S., Koonin, E. V., Cheng, D.R., and Scott, D.A. (2018). Cas13d Is a Compact RNA-Targeting Type VI CRISPR Effector Positively Modulated by a WYL-Domain-Containing Accessory Protein. Mol. Cell 70, 327–339.e5. 10.1016/j.molcel.2018.02.028.

27. O’Connell, M.R. (2019). Molecular Mechanisms of RNA Targeting by Cas13-containing Type VI CRISPR-Cas Systems. J. Mol. Biol. 431, 66–87. 10.1016/J.JMB.2018.06.029.

28. Powell, J.E., Lim, C.K.W., Krishnan, R., McCallister, T.X., Saporito-Magriña, C., Zeballos, M.A., McPheron, G.D., and Gaj, T. (2022). Targeted gene silencing in the nervous system with CRISPR-Cas13. Sci. Adv. 8. 10.1126/sciadv.abk2485.

29. Zeballos C, M.A., Moore, H.J., Smith, T.J., Powell, J.E., Ahsan, N.S., Zhang, S., and Gaj, T. (2023). Mitigating a TDP-43 proteinopathy by targeting ataxin-2 using RNA-targeting CRISPR effector proteins. Nat. Commun. 14. 10.1038/s41467-023-42147-z.

30. Morelli, K.H., Wu, Q., Gosztyla, M.L., Liu, H., Yao, M., Zhang, C., Chen, J., Marina, R.J., Lee, K., Jones, K.L., et al. (2023). An RNA-targeting CRISPR–Cas13d system alleviates disease-related phenotypes in Huntington’s disease models. Nat. Neurosci. 26, 27–38. 10.1038/s41593-022-01207-1.

31. Atilano, M.L., Grönke, S., Niccoli, T., Kempthorne, L., Hahn, O., Morón-Oset, J., Hendrich, O., Dyson, M., Adams, M.L., Hull, A., et al. (2021). Enhanced insulin signalling ameliorates c9orf72 hexanucleotide repeat expansion toxicity in drosophila. Elife 10. 10.7554/eLife.58565.

32. Wang, C., Ward, M.E., Chen, R., Liu, K., Tracy, T.E., Chen, X., Xie, M., Sohn, P.D., Ludwig, C., Meyer-Franke, A., et al. (2017). Scalable Production of iPSC-Derived Human Neurons to Identify Tau-Lowering Compounds by High-Content Screening. Stem Cell Reports 9, 1221–1233. 10.1016/j.stemcr.2017.08.019.

33. Pantazis, C.B., Yang, A., Lara, E., McDonough, J.A., Blauwendraat, C., Peng, L., Oguro, H., Kanaujiya, J., Zou, J., Sebesta, D., et al. (2022). A reference human induced pluripotent stem cell line for large-scale collaborative studies. Cell Stem Cell 29, 1685–1702.e22. 10.1016/J.STEM.2022.11.004.

34. Ai, Y., Liang, D., and Wilusz, J.E. (2022). CRISPR/Cas13 effectors have differing extents of off-target effects that limit their utility in eukaryotic cells. Nucleic Acids Res. 50, E65– E65. 10.1093/nar/gkac159.

35. Burrows, D.J., Mcgown, A., Abduljabbar, O., Castelli, L.M., Shaw, P.J., Mbbs, D., Frcp, M.D., Faan, F., Faas, F., Hautbergue, G.M., et al. (2024). RAN translation of C9ORF72-related dipeptide repeat proteins recapitulates hallmarks of motor neurone disease and identifies hypothermia as a therapeutic strategy in zebrafish. bioRxiv, 2024.01.17.576077. 10.1101/2024.01.17.576077.

36. Rihel, J., Prober, D.A., Arvanites, A., Lam, K., Zimmerman, S., Jang, S., Haggarty, S.J., Kokel, D., Rubin, L.L., Peterson, R.T., et al. (2010). Zebrafish behavioral profiling links drugs to biological targets and rest/wake regulation. Science (80-.). 327, 348–351. 10.1126/SCIENCE.1183090/SUPPL_FILE/RIHEL.SOM.PDF.

37. Kroll, F., Donnelly, J., Özcan, G.G., Mackay, E., and Rihel, J. (2023). Behavioural pharmacology predicts disrupted signalling pathways and candidate therapeutics from zebrafish mutants of Alzheimer’s disease risk genes. bioRxiv, 2023.11.28.568940. 10.1101/2023.11.28.568940.

38. Chew, J., Cook, C., Gendron, T.F., Jansen-West, K., Del Rosso, G., Daughrity, L.M., Castanedes-Casey, M., Kurti, A., Stankowski, J.N., Disney, M.D., et al. (2019). Aberrant deposition of stress granule-resident proteins linked to C9orf72-associated TDP-43 proteinopathy. Mol. Neurodegener. 14. 10.1186/s13024-019-0310-z.

39. Liu, Y., Pattamatta, A., Zu, T., Reid, T., Bardhi, O., Borchelt, D.R., Yachnis, A.T., and Ranum, L.P.W. (2016). C9orf72 BAC Mouse Model with Motor Deficits and Neurodegenerative Features of ALS/FTD. Neuron 90, 521–534. 10.1016/j.neuron.2016.04.005.

40. Hawkins, P., Morton, D.B., Burman, O., Dennison, N., Honess, P., Jennings, M., Lane, S., Middleton, V., Roughan, J. V., Wells, S., et al. (2011). A guide to defining and implementing protocols for the welfare assessment of laboratory animals: eleventh report of the BVAAWF/FRAME/RSPCA/UFAW Joint Working Group on Refinement. Lab. Anim. 45, 1–13. 10.1258/LA.2010.010031.

41. 41. Fentener Van Vlissingen, J.M., Borrens, M., Girod, A., Lelovas, P., Morrison, F., and Saavedra Torres, Y. (2015). The reporting of clinical signs in laboratory animals FELASA Working Group Report. Lab. Anim. 49, 267–283. 10.1177/0023677215584249.

42. Esanov, R., Cabrera, G.T., Andrade, N.S., Gendron, T.F., Brown, R.H., Benatar, M., Wahlestedt, C., Mueller, C., and Zeier, Z. (2017). A C9ORF72 BAC mouse model recapitulates key epigenetic perturbations of ALS/FTD. Mol. Neurodegener. 12. 10.1186/S13024-017-0185-9.

43. Boivin, M., Pfister, V., Gaucherot, A., Ruffenach, F., Negroni, L., Sellier, C., and Charlet-Berguerand, N. (2020). Reduced autophagy upon C9ORF72 loss synergizes with dipeptide repeat protein toxicity in G4C2 repeat expansion disorders. EMBO J. 39. 10.15252/EMBJ.2018100574.

44. Q, Z., J, J., TF, G., M, M.-D., L, J., A, T., S, D.G., S, G.D., MJ, R., P, K., et al. (2020). Reduced C9ORF72 function exacerbates gain of toxicity from ALS/FTD-causing repeat expansion in C9orf72. Nat. Neurosci. 23, 615–624. 10.1038/S41593-020-0619-5.

45. Charlesworth, C.T., Deshpande, P.S., Dever, D.P., Camarena, J., Lemgart, V.T., Cromer, M.K., Vakulskas, C.A., Collingwood, M.A., Zhang, L., Bode, N.M., et al. (2019). Identification of preexisting adaptive immunity to Cas9 proteins in humans. Nat. Med. 2019 252 25, 249–254. 10.1038/s41591-018-0326-x.

46. Chew, W.L., Tabebordbar, M., Cheng, J.K.W., Mali, P., Wu, E.Y., Ng, A.H.M., Zhu, K., Wagers, A.J., and Church, G.M. (2016). A multifunctional AAV-CRISPR-Cas9 and its host response. Nat. Methods 13, 868–874. 10.1038/nmeth.3993.

47. Tang, X.Z.E., Tan, S.X., Hoon, S., and Yeo, G.W. (2022). Pre-existing adaptive immunity to the RNA-editing enzyme Cas13d in humans. Nat. Med. 2022 287 28, 1372–1376. 10.1038/s41591-022-01848-6.

48. Walker, A.W., Ince, J., Duncan, S.H., Webster, L.M., Holtrop, G., Ze, X., Brown, D., Stares, M.D., Scott, P., Bergerat, A., et al. (2011). Dominant and diet-responsive groups of bacteria within the human colonic microbiota. ISME J. 5, 220–230. 10.1038/ismej.2010.118.

49. Shirley, J.L., de Jong, Y.P., Terhorst, C., and Herzog, R.W. (2020). Immune Responses to Viral Gene Therapy Vectors. Mol. Ther. 28, 709–722. 10.1016/J.YMTHE.2020.01.001.

50. Selvaraj, B.T., Livesey, M.R., Zhao, C., Gregory, J.M., James, O.T., Cleary, E.M., Chouhan, A.K., Gane, A.B., Perkins, E.M., Dando, O., et al. (2018). C9ORF72 repeat expansion causes vulnerability of motor neurons to Ca2+-permeable AMPA receptor-mediated excitotoxicity. Nat. Commun. 9, 347. 10.1038/s41467-017-02729-0.

51. Hughes, M.P., Nelvagal, H.R., Coombe-Tennant, O., Smith, D., Smith, C., Massaro, G., Poupon-Bejuit, L., Platt, F.M., and Rahim, A.A. (2023). A Novel Small NPC1 Promoter Enhances AAV-Mediated Gene Therapy in Mouse Models of Niemann–Pick Type C1 Disease. Cells 12. 10.3390/CELLS12121619/S1.

52. Fernandopulle, M.S., Prestil, R., Grunseich, C., Wang, C., Gan, L., and Ward, M.E. (2018). Transcription Factor-Mediated Differentiation of Human iPSCs into Neurons. Curr. Protoc. Cell Biol. 79, e51. 10.1002/cpcb.51.

53. Simone, R., Balendra, R., Moens, T.G., Preza, E., Wilson, K.M., Heslegrave, A., Woodling, N.S., Niccoli, T., Gilbert-Jaramillo, J., Abdelkarim, S., et al. (2018). G-quadruplex-binding small molecules ameliorate C9orf72 FTD/ALS pathology in vitro and in vivo. EMBO Mol. Med. 10, 22–31. 10.15252/emmm.201707850.

54. Licata, N. V, Cristofani, R., Salomonsson, S., Wilson, K.M., Kempthorne, L., Vaizoglu, D., D’Agostino, V.G., Pollini, D., Loffredo, R., Pancher, M., et al. (2022). C9orf72 ALS/FTD dipeptide repeat protein levels are reduced by small molecules that inhibit PKA or enhance protein degradation. EMBO J. 41. 10.15252/EMBJ.2020105026.

55. Bray, N.L., Pimentel, H., Melsted, P., and Pachter, L. (2016). Near-optimal probabilistic RNA-seq quantification. Nat. Biotechnol. 34, 525–527. 10.1038/NBT.3519.

56. Love, M.I., Huber, W., and Anders, S. (2014). Moderated estimation of fold change and dispersion for RNA-seq data with DESeq2. Genome Biol. 15. 10.1186/S13059-014-0550-8.

57. Soneson, C., Love, M.I., and Robinson, M.D. (2016). Differential analyses for RNA-seq: Transcript-level estimates improve gene-level inferences. F1000Research 4. 10.12688/F1000RESEARCH.7563.2.

58. Aleström, P., D’Angelo, L., Midtlyng, P.J., Schorderet, D.F., Schulte-Merker, S., Sohm, F., and Warner, S. (2020). Zebrafish: Housing and husbandry recommendations. Lab. Anim. 54, 213–224. 10.1177/0023677219869037.

59. Bates, D., Mächler, M., Bolker, B.M., and Walker, S.C. (2015). Fitting Linear Mixed-Effects Models Using lme4. J. Stat. Softw. 67, 1–48. 10.18637/JSS.V067.I01.

60. Häger, C., Keubler, L.M., Biernot, S., Dietrich, J., Buchheister, S., Buettner, M., and Bleich, A. (2015). Time to Integrate to Nest Test Evaluation in a Mouse DSS-Colitis Model. PLoS One 10, e0143824. 10.1371/JOURNAL.PONE.0143824.

