## Supplemental figures for "Dual-targeting CRISPR-CasRx reduces *C9orf72* ALS/FTD sense and antisense repeat RNAs in vitro and in vivo"

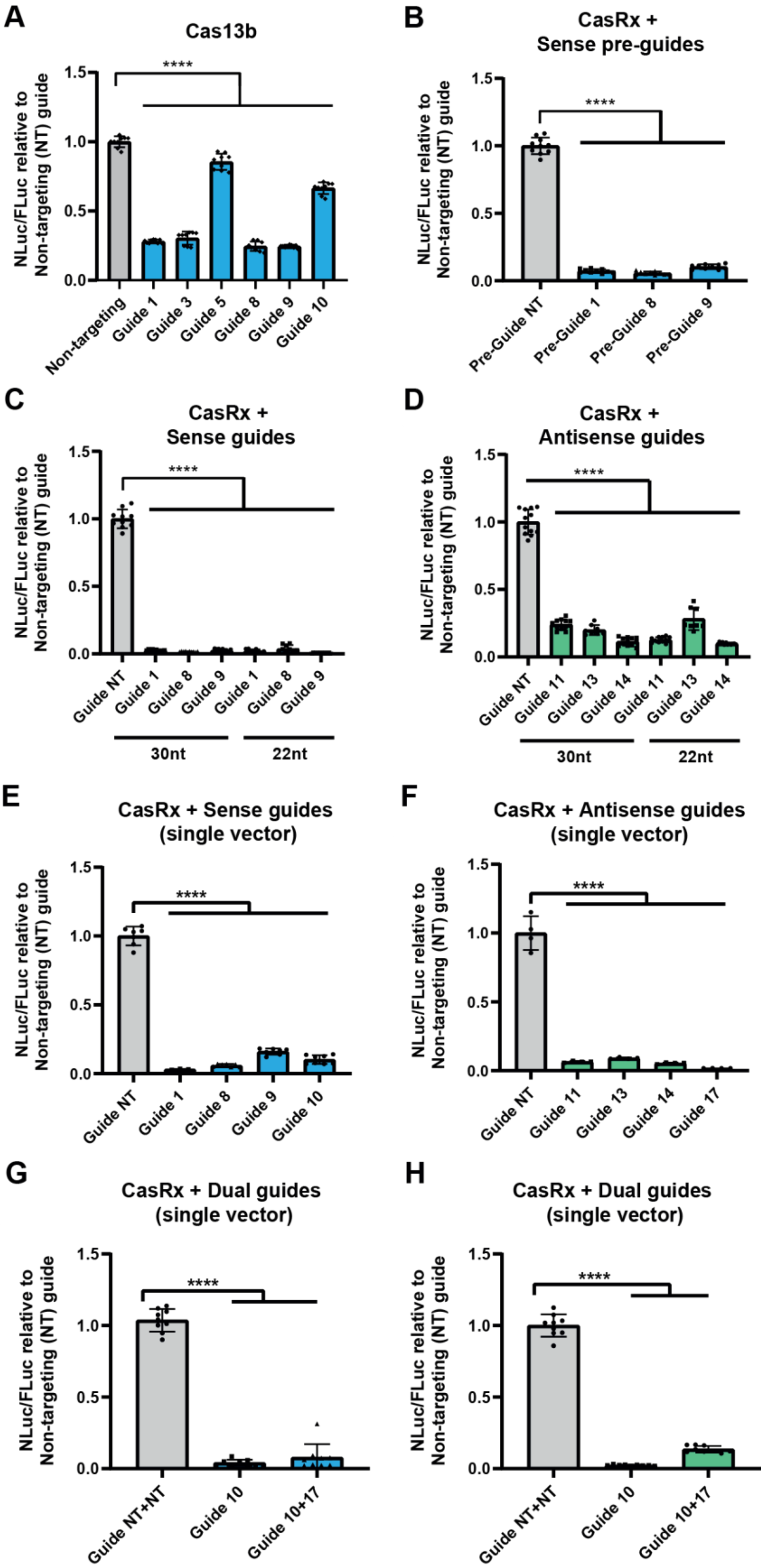
A
Cas13b
NLuc/FLuc relative to Non-targeting (NT) guide
****
Non-targeting
Guide 1
Guide 3
Guide 5
Guide 8
Guide 9
Guide 10
B
CasRx + Sense pre-guides
Pre-Guide NT
Pre-Guide 1
Pre-Guide 8
Pre-Guide 9
C
CasRx + Sense guides
Guide NT
30nt
22nt
D
CasRx + Antisense guides
Guide 11
Guide 13
Guide 14
E
CasRx + Sense guides (single vector)
F
CasRx + Antisense guides (single vector)
Guide 17
G
CasRx + Dual guides (single vector)
Guide NT+NT
Guide 10+17
H
CasRx + Dual guides (single vector)

**Figure S1. CRISPR-CasRx can mature pre-gRNAs and reduce sense and antisense repeat-containing RNAs in a single construct with higher efficacy than Cas13b. (A)** Cas13b knockdown efficiency tested with *C9orf72* sense repeat targeting guides in our sense NLuc assay. **(B)** Sense targeting guides cloned into a pre-gRNA expressing plasmid and tested in our sense NLuc assay. **(C-D)** Testing of 30 nt and 22 nt gRNA variants of previously tested gRNAs in both the **(C)** sense and **(D)** antisense NLuc reporter assays. **(E)** Sense NLuc assays testing single plasmids expressing both CasRx and sense guides. **(F)** Antisense NLuc assays testing single plasmids expressing both CasRx and antisense guides. **(G-H)** Single vectors expressing a single guide (guide 10 for sense targeting or guide 17 for antisense targeting) or both guides 10 and 17 were used in our sense and antisense NLuc reporter assays. CasRx with sense and antisense targeting array can effectively reduce both **(G)** sense and **(H)** antisense *C9orf72* DPR levels to a similar degree to single guide expressing plasmids indicating effective guide array maturation and multi-target engagement. All NLuc data normalised to FLuc and non-targeting (NT+NT) guide. Data given as mean ± S.D. n=3 biological repeats, one-way ANOVA, Holm-Sidak post-hoc analysis, ***p<0.001, ****p<0.0001.

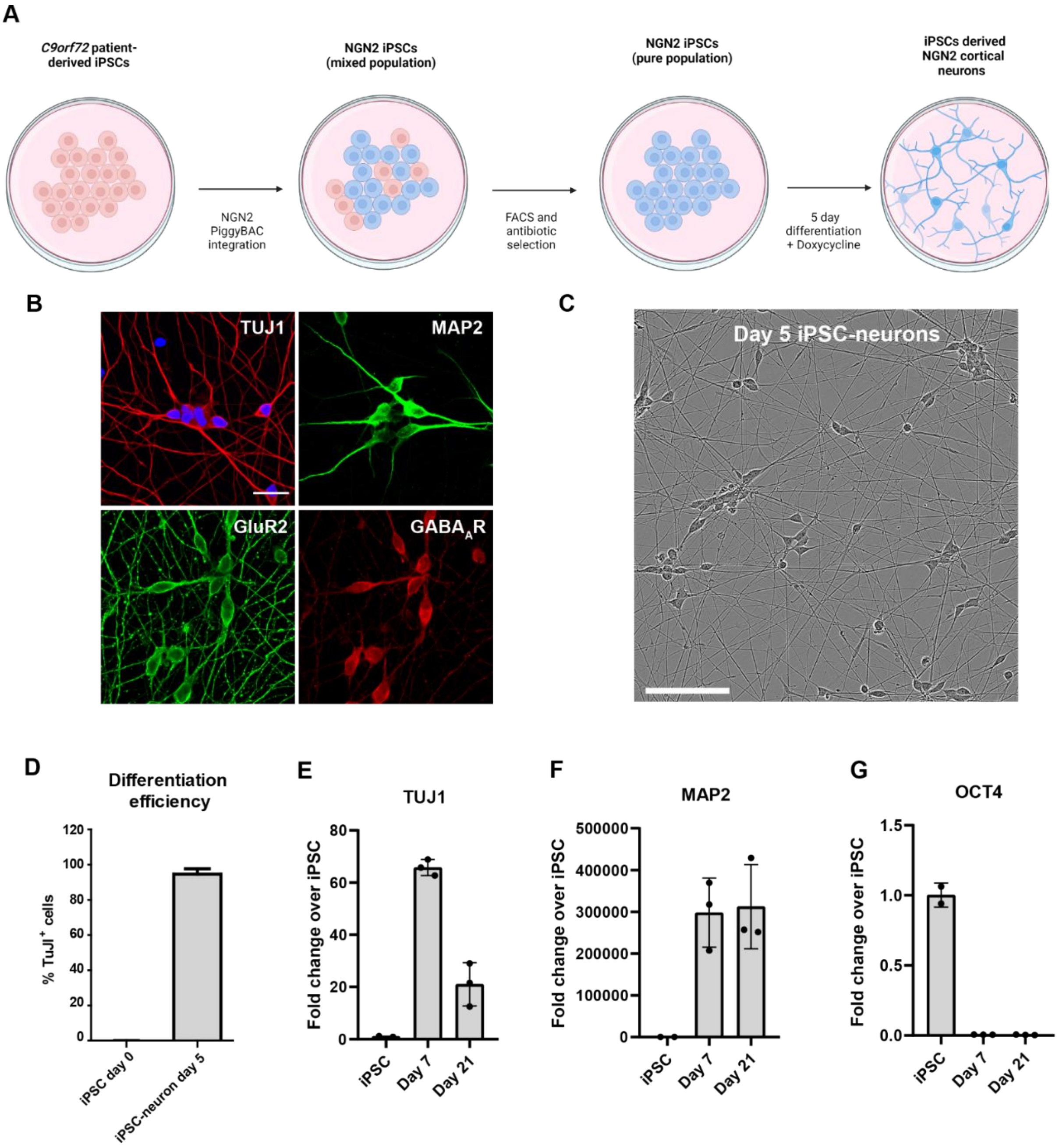
A
C9orf72 patient-derived iPSCs
NGN2 PiggyBAC integration
NGN2 iPSCs (mixed population)
FACS and antibiotic selection
NGN2 iPSCs (pure population)
5 day differentiation + Doxycycline
iPSCs derived NGN2 cortical neurons
B
TUJ1
MAP2
GluR2
GABAAR
C
Day 5 iPSC-neurons
D
Differentiation efficiency
% TuJ1+ cells
iPSC day 0
iPSC-neuron day 5
E
TUJ1
Fold change over iPSC
iPSC
Day 7
Day 21
F
MAP2
Fold change over iPSC
G
OCT4
Fold change over iPSC

**Figure S2. Characterisation of NGN2 iPSC-neuron model. (A)** Schematic NGN2 iPSC generation and $i^3$Neuron differentiation protocol. A doxycycline-inducible NGN2 + BFP cassette was genomically inserted into *C9orf72* patient-derived iPSCs with piggyBAC transposase. iPSCs were selected for presence of the cassette and then rapidly differentiated into mixed cortical-like neurons via the $i^3$Neuron protocol. **(B)** ICC of differentiated NGN2 neurons after 5 days *in vitro* (DIV5) showing expression of neuronal markers $GABA_AR$, GluR2, MAP2 and TUJ1; scale bar = 50 µm. **(C)** DIV5 NGN2 neurons morphologically resemble neurons with neuronal projections visible by light microscopy; scale bar = 200 µm. **(D)** Quantification of % of TUJ1 positive cells at DIV5. **(E-G)** RT-qPCR analysis on RNA from DIV7 and DIV21 differentiated neurons show expression of **(E)** TUJ1, **(F)** MAP2, and loss of expression of pluripotency marker **(G)** OCT4. Data presented as fold change compared to undifferentiated NGN2 iPSCs across 3 separate neuronal inductions of C9 line 1.

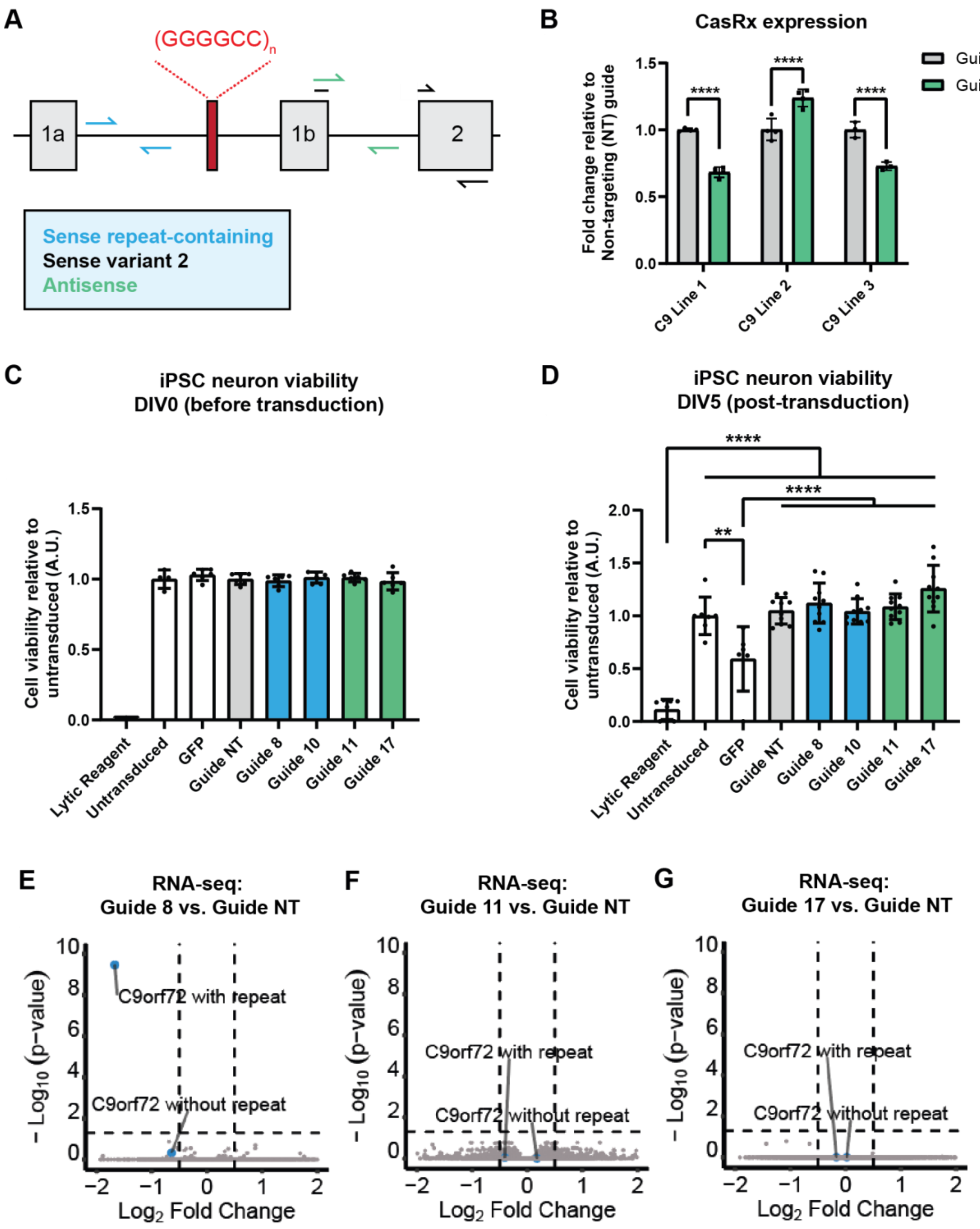
A
(GGGGCC)n
1a
1b
2
Sense repeat-containing
Sense variant 2
Antisense
B
CasRx expression
Fold change relative to Non-targeting (NT) guide
Guide NT
Guide 17
****
C9 Line 1
C9 Line 2
C9 Line 3
C
iPSC neuron viability DIV0 (before transduction)
Cell viability relative to untransduced (A.U.)
Lytic Reagent
Untransduced
GFP
Guide NT
Guide 8
Guide 10
Guide 11
Guide 17
D
iPSC neuron viability DIV5 (post-transduction)
****
**
E
RNA-seq: Guide 8 vs. Guide NT
F
RNA-seq: Guide 11 vs. Guide NT
G
RNA-seq: Guide 17 vs. Guide NT
– Log10 (p-value)
Log2 Fold Change
C9orf72 with repeat
C9orf72 without repeat

**Figure S3. CRISPR-CasRx expression in iPSC-neurons and viability/off target analysis. (A)** Schematic of RT-qPCR primers used to quantify *C9orf72* transcripts. **(B)** RT-qPCR for CasRx expression in 3 independent inductions of C9 line 1 and a single induction each of C9 lines 2 and 3, transduced with lentivirus expressing CasRx and either non-targeting (NT) guide or antisense targeting guide 17. Data presented as fold change compared to non-targeting (NT) gRNA. Data given as mean ± S.D, n=3 biological replicates of C9 line 1 and n=3 technical replicates of C9 lines 2 and 3, one-way ANOVA, Holm-Sidak post-hoc analysis, ****$p<0.0001$**. (C-D)** Cell viability of neurons treated with CRISPR-CasRx lentivirus (with sense targeting guide 8 or 10 or antisense targeting guides 11 and 17) assessed **(C)** immediately following transduction (DIV0) and **(D)** 5 days post-transduction (DIV5) via CellTiter-Fluor™ assay. Untransduced and GFP-only lentivirus treated samples were included as controls, alongside the lytic reagent, the positive assay control. Data given as mean ± S.D, n=3 biological replicates of C9 line 1, one-way ANOVA with Holm-Sidak post-hoc analysis ** $p<0.01$. ****$p<0.0001$. **(E)** Volcano plots of DESeq2 analysis showing DEGs between neurons treated with CasRx lentiviruses expressing sense-targeting guide 8 compared to CasRx lentivirus expressing non-targeting (NT) guide with *C9orf72* transcripts grouped by those that contain intron1 and the repeats, and those that do not. **(G-H)** Volcano plots of DESeq2 analysis showing no DEGs between cells treated with CasRx lentiviruses expressing antisense-targeting guides 11 **(G)** or 17 **(H)**, compared to CasRx non-targeting (NT) control lentivirus. Dotted lines indicate thresholds for fold change on x axis ($|\log_2 FoldChange|>0.5$) and *p* value on y-axis (adjusted $p<0.05$). n=3 independent inductions of C9 line 1.

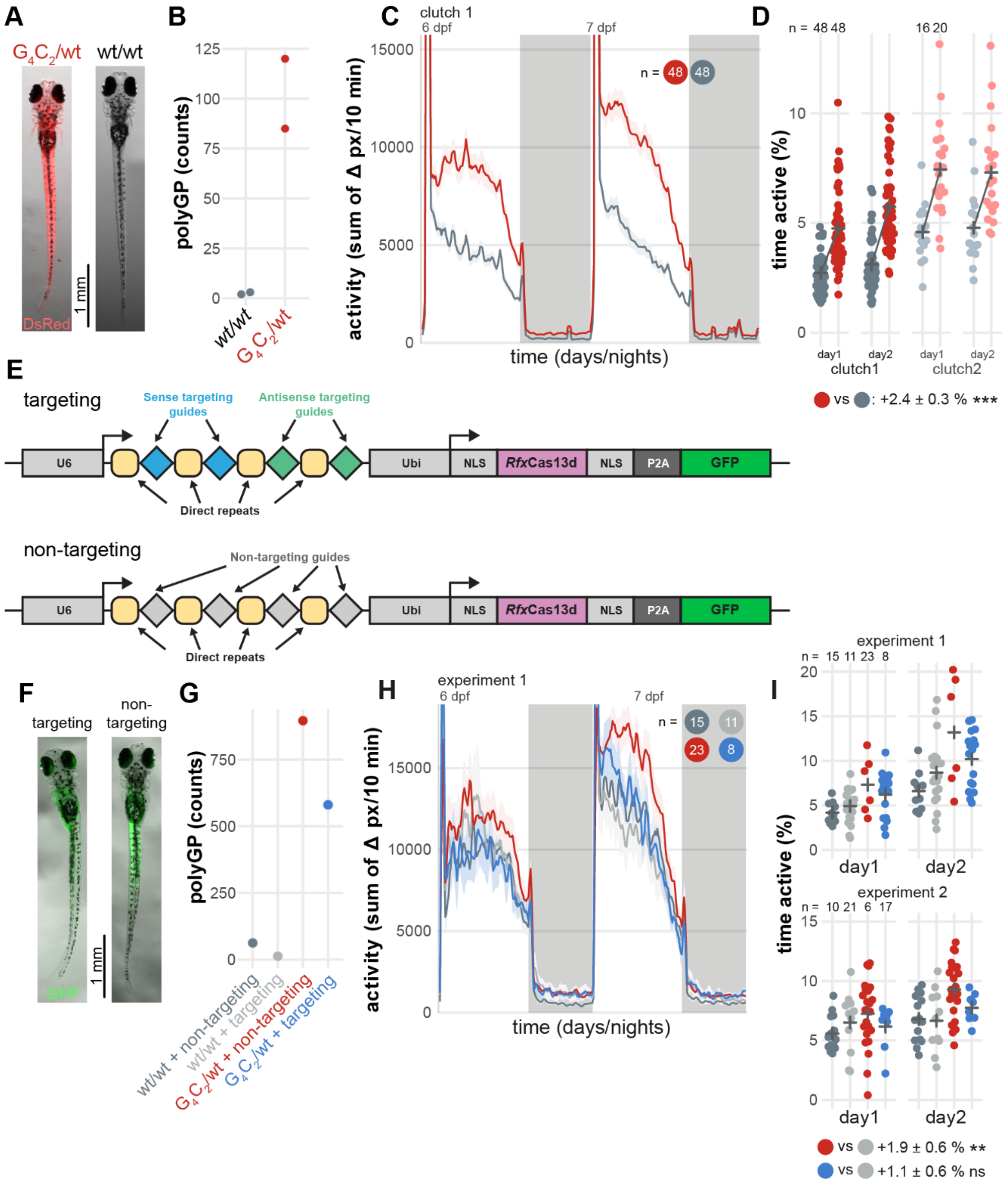
A
$G_4C_2$/wt
wt/wt
DsRed
1 mm
B
polyGP (counts)
wt/wt
$G_4C_2$/wt
C
clutch 1
6 dpf
7 dpf
n = 48 48
activity (sum of Δ px/10 min)
time (days/nights)
D
n = 48 48
16 20
time active (%)
day1
day2
clutch1
clutch2
vs : +2.4 ± 0.3 % ***
E
targeting
Sense targeting guides
Antisense targeting guides
U6
Ubi
NLS
RfxCas13d
NLS
P2A
GFP
Direct repeats
non-targeting
Non-targeting guides
U6
Ubi
NLS
RfxCas13d
NLS
P2A
GFP
Direct repeats
F
targeting
non-targeting
GFP
1 mm
G
polyGP (counts)
wt/wt + non-targeting
wt/wt + targeting
$G_4C_2$/wt + non-targeting
$G_4C_2$/wt + targeting
H
experiment 1
6 dpf
7 dpf
n = 15 11 23 8
activity (sum of Δ px/10 min)
time (days/nights)
I
experiment 1
n = 15 11 23 8
experiment 2
n = 10 21 6 17
time active (%)
day1
day2
vs +1.9 ± 0.6 % **
vs +1.1 ± 0.6 % ns

**Figure S4. CRISPR-CasRx rescues larval hyperactivity in a *C9orf72* zebrafish model. (A)** An example *ubi:$G_4C_2$×45* heterozygous larva ($G_4C_2$/wt) expressing DsRed compared to a wild-type (wt/wt) sibling at 8 dpf. **(B)** Levels of polyGP DPRs in pools of 8-dpf larvae, measured by MSD. Each dot represents one clutch. **(C)** Activity (sum of Δ pixels/10 minutes) of wild-type (wt/wt, dark grey) and $G_4C_2$ heterozygous ($G_4C_2$/wt, red) larvae during 48 hr on a 14 hr:10 hr light:dark cycle (white background for days, grey background for nights). **(D)** Time spent active (% of each day) for each larva. Black crosses mark the group means. Compared to wild-type larvae (wt/wt, dark grey), *ubi:$G_4C_2$×45* heterozygous larvae ($G_4C_2$/wt, red) spent more time active during the day (***$p<0.001$). Statistics by likelihood-ratio test on a linear mixed effect model. **(E)** Schematic of the CRISPR-CasRx plasmid injected in *ubi:$G_4C_2$×45* heterozygous larvae and wild-type embryos. The flanking Tol2 arms are not shown. NLS, nuclear localisation signal; Ubi, ubiquitin promoter. **(F)** Example GFP imaging of 8-dpf larvae expressing CasRx with either targeting or non-targeting gRNAs. Targeting or non-targeting plasmid and Tol2 recombinase mRNA were co-injected at the single-cell stage. **(G)** Levels of polyGP DPRs in pools of 8-dpf larvae, measured by MSD. Each dot represents the mean of two technical replicates. **(H)** Activity (sum of Δ pixels/10 minutes) of wild-type (wt/wt) and *ubi:$G_4C_2$×45* heterozygous ($G_4C_2$/wt) larvae expressing CasRx and targeting or non-targeting gRNAs, as in (C). **(I)** Time spent active (% of each day) for each larva. As in (D), compared to injected wild-type larvae (wt/wt + targeting, light grey), control-injected *ubi:$G_4C_2$×45* heterozygous larvae ($G_4C_2$/wt + non-targeting, red) spent more time active (**$p=0.01$). In contrast, the activity of *ubi:$G_4C_2$×45* heterozygous larvae expressing CasRx and targeting gRNAs ($G_4C_2$/wt + targeting, blue) was not significantly different to injected wild-type larvae (light grey). Statistics by likelihood-ratio test on a linear mixed effect model.

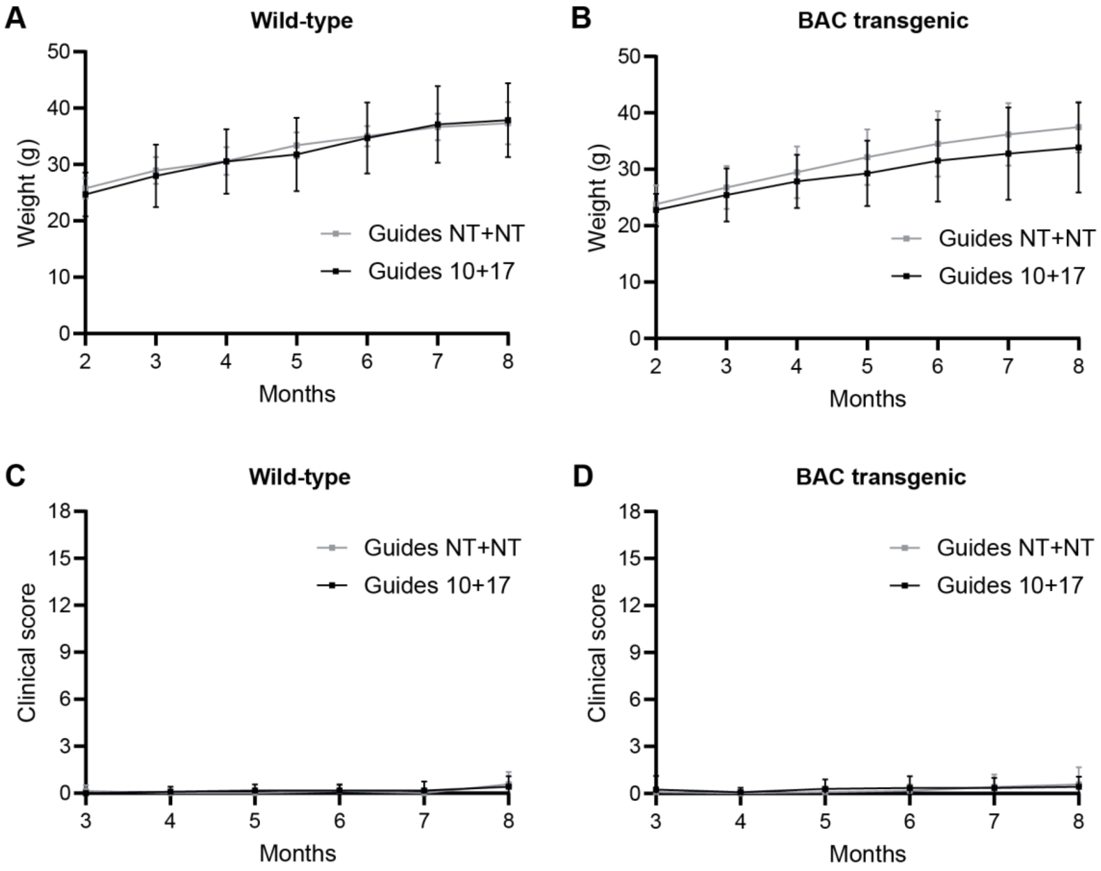
A
Wild-type
Weight (g)
Months
Guides NT+NT
Guides 10+17
B
BAC transgenic
Weight (g)
Months
Guides NT+NT
Guides 10+17
C
Wild-type
Clinical score
Months
Guides NT+NT
Guides 10+17
D
BAC transgenic
Clinical score
Months
Guides NT+NT
Guides 10+17

**Figure S5. CRISPR-CasRx treatment does not cause overt toxicity *in vivo* in wild-type or *C9orf72* BAC transgenic mice. (A-B)** Monthly body weight analysis of **(A)** wild-type or **(B)** *C9orf72* BAC transgenic mice injected at P0 with either non-targeting (Guides NT+NT) or dual sense and antisense targeting (Guides 10+17) CasRx PHP.eB AAV from 2 months of age until end of the study. **(C-D)** Composite clinical score of **(C)** wild-type or **(D)** *C9orf72* BAC transgenic mice injected at P0 with either non-targeting or Guide 10+17 CasRx PHP.eB AAV from 2 months of age until end of the study. Wild-type: n=12 Guide 10+17; n=7 Guide NT+NT. *C9orf72* BAC: n=14 Guide 10+17; n=12 Guide NT+NT.
